# Pathological tau activates inflammatory nuclear factor-kappa B (NF-κB) and pT181-Qβ vaccine attenuates NF-κB in PS19 tauopathy mice

**DOI:** 10.1101/2025.03.10.642500

**Authors:** Karthikeyan Tangavelou, Shanya Jiang, Somayeh Dadras, Jonathan P Hulse, Kathryn Sanchez, Virginie Bondu, Zachary Villaseñor, Michael Mandell, Julianne Peabody, Bryce Chackerian, Kiran Bhaskar

## Abstract

Tau regulates neuronal integrity. In tauopathy, phosphorylated tau detaches from microtubules and aggregates, and is released into the extracellular space. Microglia are the first responders to the extracellular tau, a danger/damage-associated molecular pattern (DAMP), which can be cleared by proteostasis and activate innate immune response gene expression by nuclear factor-kappa B (NF-κB). However, longitudinal NF-κB activation in tauopathies and whether pathological tau (pTau) contributes to NF-κB activity is unknown. Here, we tau oligomers from human Alzheimer’s disease brain (AD-TO) activate NF-κB in mouse microglia and macrophages reducing the IκBα via promoting its secretion in the extracellular space. NF-κB activity peaks at 9- and 11-months age in PS19Luc^+^ and hTauLuc^+^ mice, respectively. Reducing pTau via pharmacological (DOX), genetic (*Mapt^-/-^*) or antibody-mediated neutralization (immunization with pT181-Qβ vaccine) reduces NF-κB activity, and together suggest pTau is a driver of NF-κB and chronic neuroinflammation tauopathies.

**Summary:** Neuronal tau activates microglial NF-κB constitutively by secreting its inhibitor IκBα. NF-κB activation in PS19Luc^+^ and hTauLuc^+^ mice peaks at 9- and 11-months of age, respectively. Neutralizing pTau with pT181-Qβ vaccine (targeting phosphorylated threonine 181 tau) alleviates NF-κB activity in tauopathy mice.

## Introduction

Human *MAPT* gene expresses six isoforms of tau (2N4R, 1N4R, 0N4R, 2N3R, 1N3R, 0N3R) in the brain by alternative splicing of exons^1^. Tauopathy is a group of more than 20 neurodegenerative diseases caused by aggregation of hyperphosphorylated tau^2,3^. Alzheimer’s disease (AD) is one of the tauopathies and its brain pathology is characterized as extracellular aggregation of amyloid beta peptide (Aβ) and intracellular accumulation of hyperphosphorylated tau as neurofibrillary tangles (NFTs)^4^. There are many factors which can cause tau phosphorylation and aggregation, including excessive cytokines secretion from microglia^5^, viral^6,7^ or bacterial infections^8,9^, bacterially derived lipopolysaccharide (LPS)^10^, Aβ aggregates^11^, oxidative stress ^12^, unfolded protein response (UPR) ^13^, inherited or sporadic mutations ^14^, environmental toxins ^15^, aging ^16,17^, sleep apnea ^18,19^, and drug and alcohol abuses ^20,21,22,23,24,25^. Aggregation of tau indicates dysfunctional proteostasis in neurons, which are postmitotic cells and cannot undergo cell division to dilute protein aggregates or restore proteostasis function to clear protein aggregates^26^. However, neurons can transmit abnormal tau from neurons to neurons or extracellular space to dilute tau toxicity as part of cell survival mechanisms^27^. Since the abnormal tau is seeding competent, it can propagate tau pathology from neurons to neurons, which provides a tau source for seeding aggregates in healthy neurons ^28,29^. The brain resident macrophage-like immune cells, microglia, are the first responders to the extracellular tau ^30,31^ Microglia consider tau as danger/damage-associated molecular pattern (DAMP) to endocytose or phagocytose to clear tau by proteostasis and also activates the innate immune response gene expression by nuclear factor-kappa B (NF-κB) against the invader tau^5,32,33^.

While the transcription factor, NF-κB was initially identified as an immune response gene activator in B cells^34^, the subsequent findings revealed that NF-κB is a ubiquitously expressed factor in eukaryotes and regulates the gene expression associated with cell survival and apoptosis^35,36^ The endogenous inhibitor of NF-κB is IκB isoforms, which sequester the dimers of NF-κB (p50 (NF-κB1)-p65 (NF-κB3/RELA)/c-Rel) as an inactive form in the cytoplasm. Degradation of IκB allows the active NF-κB dimers to enter into the nucleus, where NF-κB binds to the promoter region of DNA and activates the gene expression^37,38^. IκBα, is the most studied isoform of NF-κB inhibitor, has been shown as a substrate of 20S proteasome^39^, 26S proteasome^40^, autophagy^41^, chaperone-mediated autophagy (CMA)^42^, and calpain^43,44^ depending on the contexts such as cell types, stimuli, and acute or chronic exposure with various stimuli. However, the tau-mediated IκBα degradation mechanism for NF-κB activation in microglia is unknown.

NF-κB activity in neurons is beneficial by positively regulating neurite outgrowth ^45,46^ synaptogenesis, plasticity ^47,48^ and long-term memory formation ^49,50^. NF-κB activation in microglia by extracellular pathological tau is detrimental and spreads tau pathology in neurons by microglial-derived interleukin-1β (IL-1β)^5,51^. Strikingly, our prior study suggested that deficiency of inflammatory adaptor signaling proteins, myeloid differentiation primary response 88 (MyD88) or apoptosis-associated speck-like protein containing a CARD (ASC) in microglia decreases tau-induced NF-κB activation, IL-1β maturation, and cognitive impairment in a hTau tauopathy mouse model^5^. Reactive microglia are known to process seeding competent tau and propagate tau pathology in neurons^52,53^. The inhibitory κB kinase (IKK) complex consists of IKKα-IKKβ-IKKγ, which regulates IκBα degradation for NF-κB activation and IKKγ/NEMO recruits autophagy receptors for protein aggregates clearance ^54,55^. Knocking out the *Ikkb* gene in microglia inhibits NF-κB activation. It restores lysosomal activity such as CMA in microglia, which consequently prevents spreading of seeding competent tau in neurons and improves spatial memory in a tauopathy PS19 mouse model^34^. However, inhibiting NF-κB activity in microglia does not clear tau aggregates in PS19 tauopathy mice brains^34^. It is therefore essential to develop an effective therapeutic strategy to clear tau aggregates and prevent excessive NF-κB activation in microglia.

Pathological tau, such as phosphorylated tau at threonine T181 (pT181), a phosphorylated tau, has been widely used as a standard and earliest biomarker at blood plasma and cerebrospinal fluid (CSF) levels for AD^56,57,58,59^. Our previous study developed a vaccine against p-Tau T181 using a viral-like particle (VLP) from bacteriophage Qbeta RNA virus backbone^60^. A 13 amino acid residues antigenic peptide from p-Tau T181 region was covalently conjugated with VLP-Qß (pT181-Qß) and immunized into tauopathy rTg4510 transgenic mice carrying frontotemporal dementia (FTD)-associated human Tau P301L mutant^64^. Interestingly, pT181-Qß vaccinated rTg4510 mice decreased microgliosis and tau pathology, including phosphorylated tau and NFTs levels, and improved spatial memory deficits^64^. However, it is unknown if pT181-Qß vaccine can prevent pathological tau-mediated NF-κB activation, which exacerbates tau pathology in AD.

This study shows purified tau monomer, oligomers, and paired-helical filaments (PHFs) induce NF-κB-GFP-Luciferase (NGL) activity in mouse BV2 microglia and bone marrow-derived macrophages (BMM). To understand the mechanism of endogenous NF-κB inhibitor IκBα degradation for NF-κB activation in C20 human microglia line, we induced NF-κB activity with tau oligomers isolated (AD-TO) from autopsy of human AD brains and identified that NF-κB is constitutively activated by secreting IκBα into the conditioned media. To assess the longitudinal effects of pathological tau in NF-κB activation, NGL albino mice were crossed with tauopathy black mice of PS19 and hTau, respectively. Our in vivo data show that NF-κB luciferase activity begins at 6-, and peaks at 9- and 11-months aged tauopathy albino mice PS19Luc^+^ and hTauLuc^+^, respectively. Tau may function as a DAMP sensor and exacerbate neuroinflammation with LPS in NGL mice, but not *Mapt^-/-^*Luc^+^ mice. Our Qβ-pT181-vaccine alleviated tau pathology and NF-κB activity in PS19Luc^+^ mice.

## Results

### Pathological tau stimulates inflammatory NF-κB activity in myeloid cells

To address the direct role of extracellular pathological tau in inflammatory NF-κB activation in myeloid cells, we used purified tau monomer, oligomers and neurofibrillary tangles as inflammatory stimuli that were isolated from autopsy of frontotemporal lobar degeneration (FTLD) or Braack stage VI of AD human brains (kindly shared by Dr. Rakez Kayed at the University of Texas Medical Branch). The resulting purified pathological tau structure and morphology were resolved at nanoscale resolution using a super-resolution direct stochastic optical reconstruction microscopy (dSTORM) imaging of tau monomer and oligomers, and transmission electron microscopy visualization of neurofibrillary tangles such as straight and paired helical filaments (Fig. 1A). Human brain-derived monomeric and oligomeric tau was spotted on the coverslip and stained with the human tau antibody, Tau12. We observed tau oligomers appeared more organized than tau monomers with larger aggregates of monomeric species several of them were larger than 10nm in diameter (Fig.1A). Next, we performed transmission electron microscopy (TEM) analysis of Sarkosyl insoluble assay-based purification of straight filaments (SFs) and paired helical filaments (PHFs) purified from autopsy of FTLD-associated tauopathy human brains. As expected, several >50nm longer SFs and PHFs were detectable from the tauopathy FTLD brains with PHFs showing clear twisted ribbon-like morphology.

**Fig. 1.**
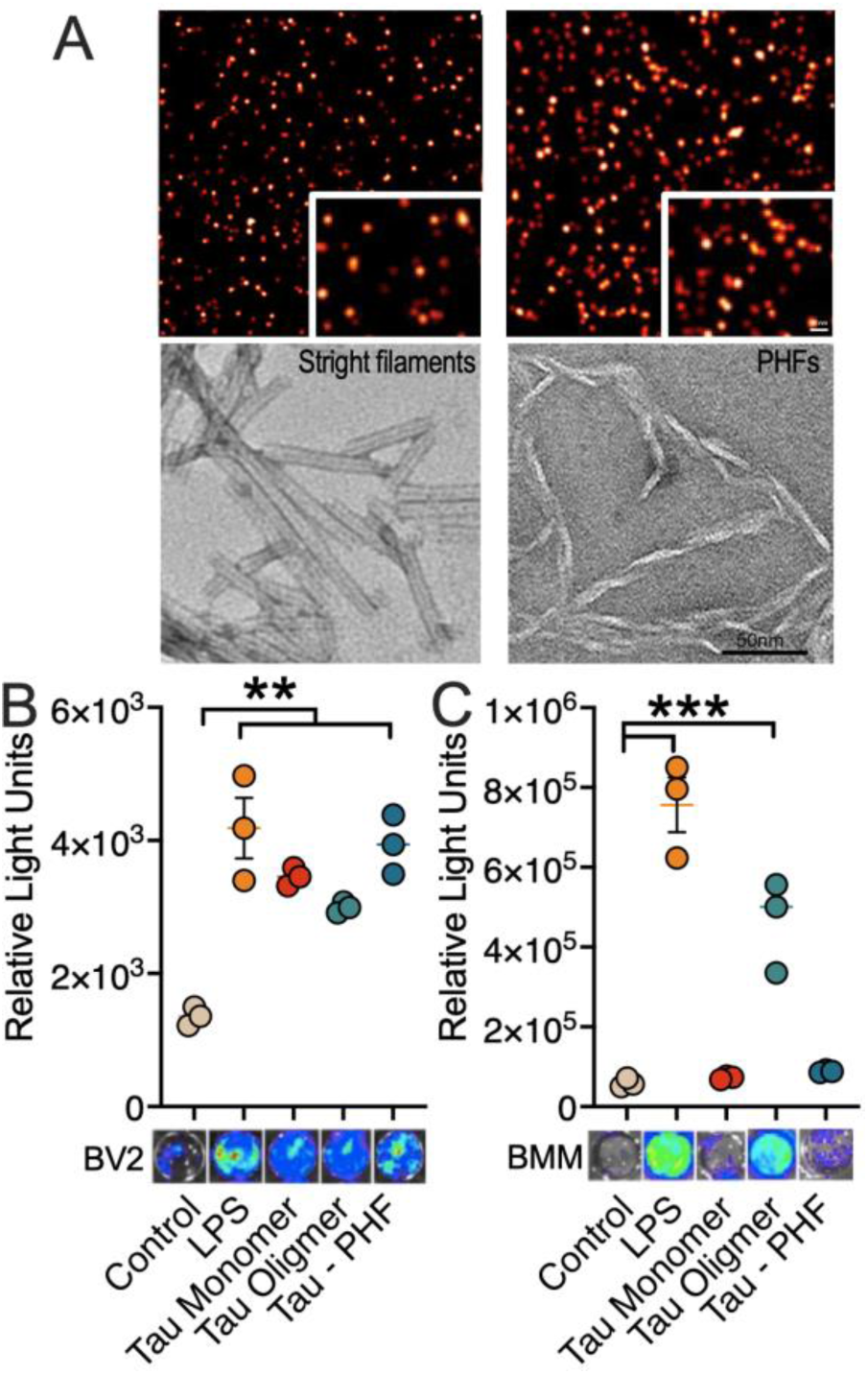
Bioluminescence analysis suggest human brain-derived pTau induces NF-kB activation in human primary microglia. **A.** Super-resolution direct stochastical optical reconstruction microscopy (dSTORM) imaging (top two panels) and electron-microscopy of size-exclusion chromatography purified tau monomers, tau oligomers, NFTs (straight and paired helical filaments) from human FTLD-Tau brains. Scale 10nm (inset) and 50 nm. **B.** BV2 cells (1x105 cells/well) transduced with lentiviral particle encoding NGL reporter were treated with LPS (200 ng/ml; 12 h) show robust NF-kB activation (left and middle histograms). BV2 cells treated with human FTLD brain-derived tau monomers, oligomers or PHFs (2 mg/ml; 12 h), showed NF-kB activation upon luciferin treatment compared to controls (vehicle). **C.** Bone-marrow derived macrophages (BMM) from NGL mice stimulated with LPS, tau monomers, tau oligomer and PHF show NF-kB activation in LPS and tau oligomer groups and less in other groups. Data displayed as mean + SEM, unpaired t test or one-way ANOVA, *p<0.05, **p<0.01; ***p < 0.005 with n=3).

We used mouse BV2 microglia cell line expressing NF-κB-GFP-Luciferase (NGL) and primary macrophages isolated from bone marrow of NGL reporter mice as source of myeloid cells to address pathological tau stimulate NF-κB activity. We have previously demonstrated that human brain-derived PHFs induce many NF-κB-related genes in primary microglial cells in the human brain via bulk RNA sequence analyses^5^. The gram-negative bacterial cell surface lipopolysaccharide (LPS) as a known inflammatory NF-κB stimulus was used as a positive control for comparative analysis of active NF-κB luciferase, which was detected using the D-luciferin substrate. To directly assess NF-κB activation by tau monomer or pathological tau forms, including AD-brain derived tau oligomers (AD-TO) and FTLD-brain derived SFs/PHFs. We performed a series of NF-κB activation reporter assays using lentiviral particles encoding NF-κB-GFP-Luciferase (NGL) plasmid in BV2 microglia. As expected, lipopolysaccharide (LPS) treatment (200 ng/ml; for 12 h) significantly elevated NF-κB activity in BV2 cells (Fig. 1B). We directly treated BV2 cells expressing NGL reporter with tau monomer, oligomers and PHFs derived from human post-mortem AD/FTLD tauopathy brains and observed a significant increase in NF-κB luciferase in BV2 cells, but not with vehicle treated cells. (Fig.1B). To confirm if pathological tau induces NF-κB activation in primary myeloid cells, we isolated bone marrow-derived macrophages (BMMs) from the transgenic NGL reporter mice. We treated BMMs with tau monomer, oligomers, and PHFs 2mg/ml for 12h. Interestingly, only tau oligomers induced robust NF-κB activation that was comparable to the response from LPS (Fig. 1C). Tau monomer and PHFs showed NF-κB activation at much lower level than tau oligomers, yet were slightly higher than controls (Fig.1C). In contrast to BV2 microglia, BMMs responded more vigorous NF-κB luciferase activity to tau oligomers, which are known to be more pathogenic and seeding competent than tau monomer and PHFs (Fig. 1C). Thus, our in vitro study confirms that the extracellular pathological tau stimulates inflammatory NF-κB in myeloid cells such as microglia and macrophages, which are pro-inflammatory and accelerates tau pathology in the brains.

### Tau oligomers from AD brain (AD-TO)-induced NF-κB activation occur via sustained degradation of inhibitor of κBα (IκBα)

The rate-limiting step in activating NF-κB is the requirement of the degradation of IκBα, which otherwise stays bound to NF-κB and prevents it from entering the nucleus. We have previously demonstrated nuclear localization of p65 in BMMs treated with human PHFs via Cellomics high-content automated microscopy^5^. Since tau oligomers are known to be highly pathogenic, seeding competent, and stimulated NF-κB stronger than tau monomer or PHFs in mouse BMM (Fig. 1C), we decided to use AD-TO to stimulate NF-κB and determine the mechanism of IκBα degradation in human C20 microglia cell line. C20 cells were chronically treated with AD-TO (1mg/ml) for 48h and immunoblotted to quantify the levels of NF-κB activation and IκBα degradation. Western blot analysis revealed that C20 microglia endocytosed the extracellular AD-TO from growth media and accumulated as detected by Tau12, but not in vehicle treated cells (Fig. 2A, B). Interestingly, our western blot data shows that AD-TO increases the level of p-P65 S536 (NF-κB3) without changing the level of IκBα (Fig. 2A, C, D), which indicates active NF-κB derives the expression of IκBα. Since IκBα synthesis and degradation are dynamic metabolic processes, we blocked the protein translation with 50µM cycloheximide during NF-κB stimulation with AD-TO and 5h before harvesting the cells. Thus, obtained western blots data confirms that AD-TO induces increased level of phosphorylated p-P65 S536, an active NF-κB (Fig. 2C, 3D) and decreases the levels of NF-κB inhibitor protein, IκBα upon adding CHX (Fig. 2E, 3F). However, the mechanism of IκBα degradation for NF-κB activation in AD-TO treated microglia is unknown.

**Figure 2.**
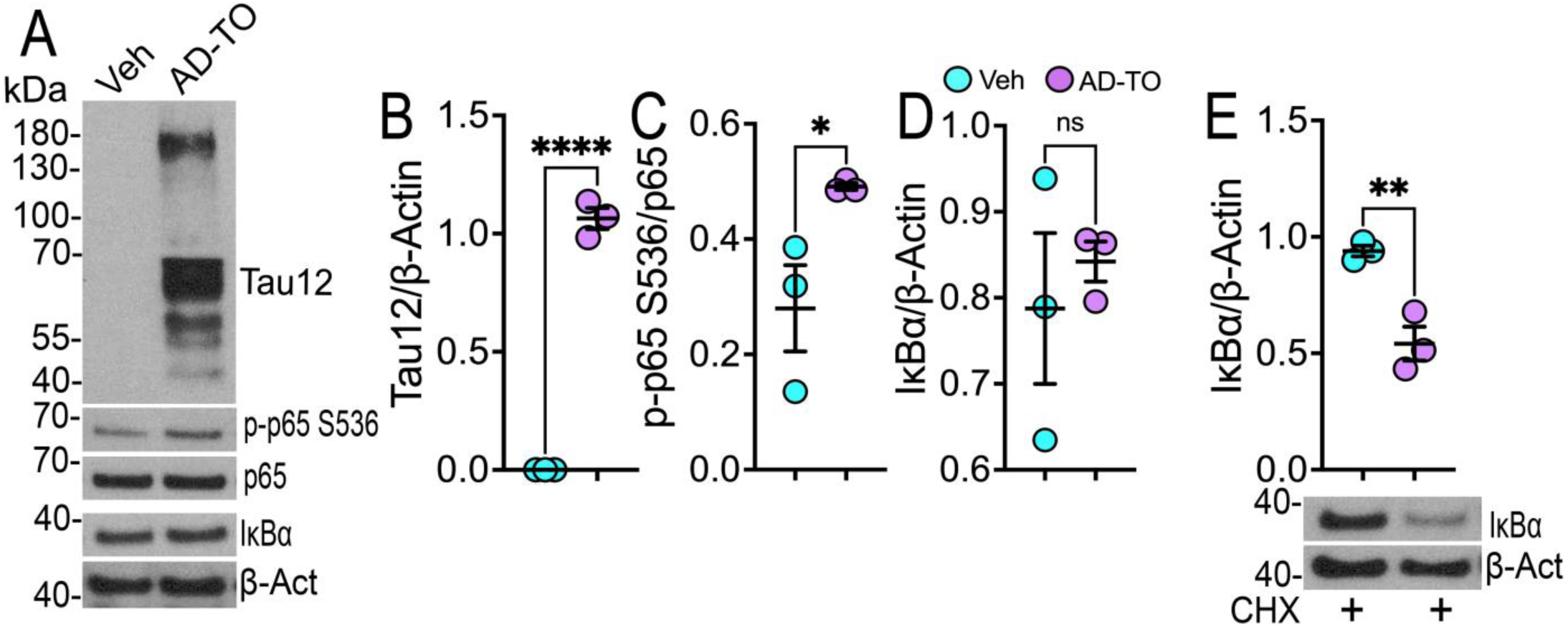
Alzheimer’s disease-tau oligomers (AD-TO) activates NF-κB by degrading its endogenous inhibitor, IκBα. **(A-E)** Human microglia C20 cells were treated with AD-TO (1mg/ml, 48h) and immunoblotted using various antibodies to quantify the levels of proteins using β-Actin as a loading control (A). AD-TO accumulated in microglia (B), which correlates with elevated phosphorylated NF-κB3 (p-P65 S536) (C) without changing the level of the endogenous NF-κB inhibitor, IκBα (D). Since IκBα level is a dynamic process associated with NF-κB-mediated synthesis and degradation, protein translation was inhibited with 50µM cycloheximide (CHX) for 5h prior harvesting the cells for western blot quantification of IκBα. Decreased level of IκBα indicates AD-TO induces NF-κB activation by degrading its endogenous inhibitor IκBα by unknown mechanism (E). Data represents as mean + SEM, unpaired t test, *p<0.05, **p<0.01; ****p < 0.005 with n=3.

### AD-TO activates NF-κB in microglia by secreting its inhibitor, IκBα into the conditioned media

Prior studies have demonstrated that IκBα is a proteasome, autophagy and calpain substrate. Upon inflammatory stimuli or infection, the primary mode of NF-κB activation occurs via IκBα degradation by one of these three major pathways. To determine the mechanism of IκBα degradation in AD-TO treated microglia, we used various proteostasis inhibitors in the presence of protein translation inhibitor, CHX. During NF-κB stimulation with AD-TO (1mg/ml) chronically for 48h, 50µM CHX (C) was added at 43h and an hour later proteostasis inhibitors were added at 44h and continued treatment until harvesting the cells at 48h for western blotting. The 20S proteasome, 26S proteasome, autophagy and calpains were inhibited with 10µM lactacystin, 10µM MG132, 100nM Baf A1 and 100µM PD 150606, respectively. Surprisingly, none of the proteostasis inhibitors individually prevented degradation of IκBα (Fig. 3A, B, F). Inhibiting calpain with PD 1505606 might have restored autophagy, activating degradation of IκBα. Interestingly, tau accumulation was significantly decreased in microglia treated with PD 150606 (Fig. 3A, C), which supports our hypothesis that inhibiting calpain might have restored tau or IκBα autophagy degradation. We therefore decided to treat microglia with the combination of MG132 (26S proteasome), Baf A1 (autophagy) and PD 150606 (calpain 1/2). Again, none of the proteostasis inhibitors either individually (Fig. 3A, B, F) or combination of all prevented IκBα degradation (Fig. 3E). The active NF-κB, phosphorylated p-P65 S536 level also significantly increased in the presence of proteostasis inhibitors except with MG132, which also showed increased level of active NF-κB but not significantly (Fig. 3D).

**Figure 3.**
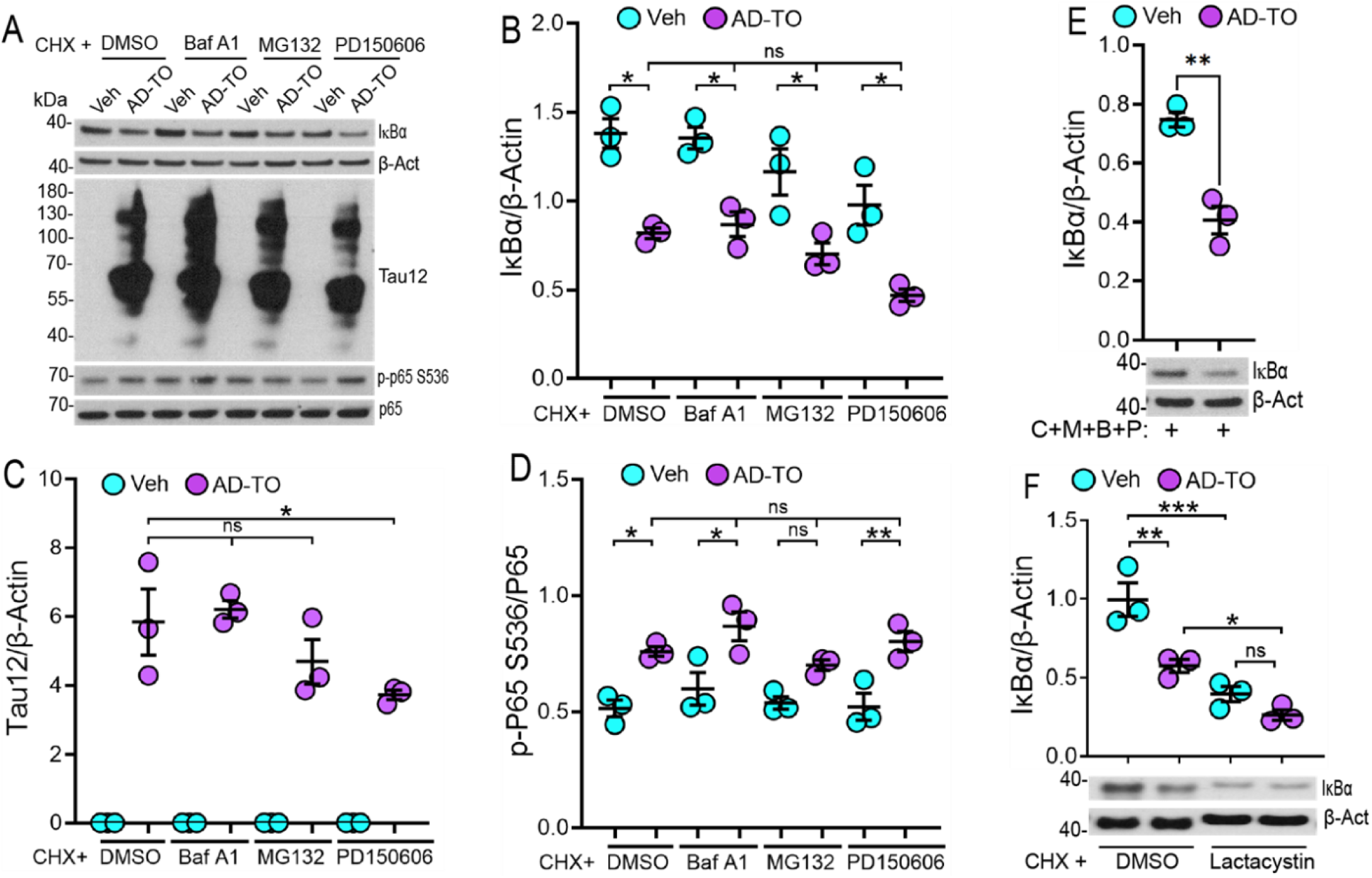
Inhibitors of proteasome, autophagy and calpain did not prevent NF-κB activation and IκBα degradation in AD-TO treated microglia. **(A-F)** Human microglia C20 cells were treated with AD-TO (1mg/ml, 48h) with or without proteostasis inhibitors and quantified the levels of active NF-κB3 and its inhibitor IκBα (A). During stimulating C20 cells with AD-TO chronically for 48h, 50µM CHX (C) was added at 43h to inhibit the protein translation and an hour later at 44h, cells were treated with proteostasis inhibitors until harvesting the cells. The 20S proteasome, 26S proteasome, autophagy and calpains were inhibited with 10µM lactacystin (L), 10µM MG132 (M), 100nM Baf A1 (B) and 100µM PD 150606 (P), respectively. Western blot quantification shows none of the proteostasis inhibitors prevent IκBα degradation (B, E, F). Tau accumulation was significantly decreased in calpain inhibitor treated microglia as compared to inhibiting the 26S proteasome or autophagy (C). Decreased level of IκBα is positively correlating with increased NF-κB activity as reflected in the level of p-P65 S536 except with MG132, which is not significantly increased (D). Data are shown as mean + SEM, unpaired t test or one-way ANOVA, *p<0.05, **p<0.01; ***p < 0.005 with n=3.

Since none of proteostasis inhibitors prevented IκBα degradation intracellularly in microglia, we searched for the presence of IκBα in conditioned media (CM). Thus, obtained CM from AD-TO+C+M+B+P treated microglia was differentially centrifuged (Fig. 4A) to fractionate small extracellular vesicles (SEV) and large extracellular vesicles (LEV) for immunoblotting of IκBα. Surprisingly, the LEV fraction obtained at 10,000g shows the presence of higher molecular weight of IκBα, which is most likely polyubiquitinated (Fig. 4B). IκBα was not detectable in the SEV fraction obtained at 100,000g, suggesting IκBα does not secrete through exosomes. Moreover, posttranslational modified IκBα was not detectable in whole cell lysates. Still, supernatant fraction obtained at 100,000g centrifugal speeds shows the presence of multiple molecular weights of IκBα, which are most likely unmodified, phosphorylated and polyubiquitinated IκBα (Fig. 4B). Together, these results suggest that AD-TO induced IκBα degradation appears to occur independent of canonical protein turnover pathways and in conditions where all three major proteostatic pathways are inhibited, it triggers secretary pathway by which polyubiquitinated IκBα is secreted into the LEVs, most likely by secretory autophagy^61^. Thus, AD-TO activates NF-κB by secreting its inhibitor IκBα upon proteostasis impairment in microglia. It is a novel finding and will be our future directions to understand how pathological tau impairs proteostasis activity and activates NF-κB in microglia whose dysfunction exacerbates neuroinflammation in AD and tauopathy brains.

**Figure 4.**
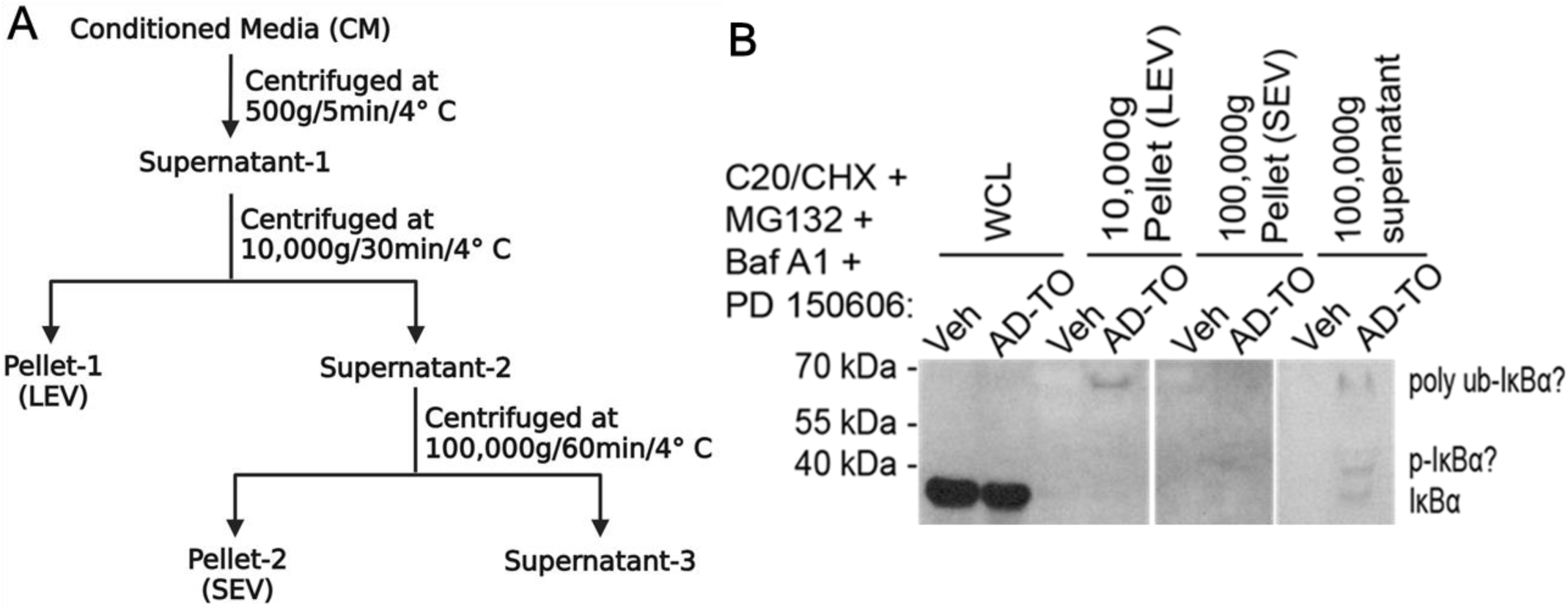
AD-TO induces secretion of NF-κB inhibitor, IκBα into the conditioned media. **(A)** A schematics of differential centrifugation steps to isolate large extracellular vesicles (LEV) and small extracellular vesicles (SEV) using the conditioned media (CM) of AD-TO treated C20 microglia. **(B)** During stimulation of C20 cells with AD-TO chronically for 48h, 50µM CHX and proteostasis inhibitors were added at 43h and 44h, respectively. CM was collected to isolate SEV and LEV (A), whereas cell pellets were processed for western blot analysis (B). Representative western blot shows the presence of IκBα in whole cell lysate (WCL) in which posttranslational modified IκBα was not detectable at longer exposure. LEV fraction shows the presence of higher molecular weight of IκBα, which is most likely polyubiquitinated, whereas IκBα was not detectable in SEV fraction. AD-TO upon impairment of proteostasis activity in microglia most likely activates the NF-κB by secreting its endogenous inhibitor IκBα into the extracellular space (B).

### NF-κB activation in the head region of PS19Luc^+^ mice is elevated as early as 6-months of age, and significantly higher at 9-months of age

To develop a model where we can longitudinally determine NF-κB activation in vivo in the head-region and the whole body, we first tested whether or not we could detect bioluminescence signal upon NF-κB activation in non-transgenic C57Bl/6j mice with black fur and NF-κB-GFP-Luciferase (NGL or Luc^+^) reporter mice with white fur in FVB genetic background, where the expression of GFP and luciferase is driven by the NF-κB promoter. We injected a small group of mice with lipopolysaccharide (LPS) (5 mg/kg; intraperitoneal, single dose). After 3h, the mice were injected with D-luciferin and subjected to bioluminescence imaging in the IVIS^®^ Spectrum advanced preclinical optical imaging system, which combines high throughput and complete tomographic optical imaging in one platform. While the Luc^+^ mice with white fur showed robust, systemic bioluminescence signal indicating the NF-κB activation by LPS, C57Bl/6j mice with black fur showed no bioluminescence (data not shown). Even removing the fur and imaging C57Bl/6j mice did not show any bioluminescence signal in response to LPS (not shown). This preliminary assessment suggested maintaining the target mice in an albino background to obtain reliable bioluminescence signal for longitudinal imaging.

To generate PS19Luc^+^ mice, we crossed Luc^+^ mice with PS19 mouse model of tauopathy (Fig. 5A). The original PS19 was in a C57Bl/6j background with black fur color. On the other hand, the NGL/Luc mice were in the FVB background, which were albino. It took over five generations of backcrossing and genotyping until we obtained PS19Luc^+^ mice with white fur (Fig. 5A). Next, we confirmed the PS19Luc^+^ mice express human P301S tau transgene by genotyping (example is shown in Supplemental Figure 5D) and determining the levels of tau hyperphosphorylation in both original PS19 and PS19Luc^+^ mice. Like PS19 mice, hippocampal lysate from PS19Luc^+^ mice showed significantly increased AT8^+^ and AT180^+^ hyperphosphorylated and Tau12^+^ total human tau (Supplemental Fig. 1A-B). Notably, there was a 36% reduction of hippocampal volume in the MRI T2 weighted analyses (Supplemental Fig. 1C), similar to those reported for original PS19 mice as reported previously^71,84^.

**Fig. 5.**
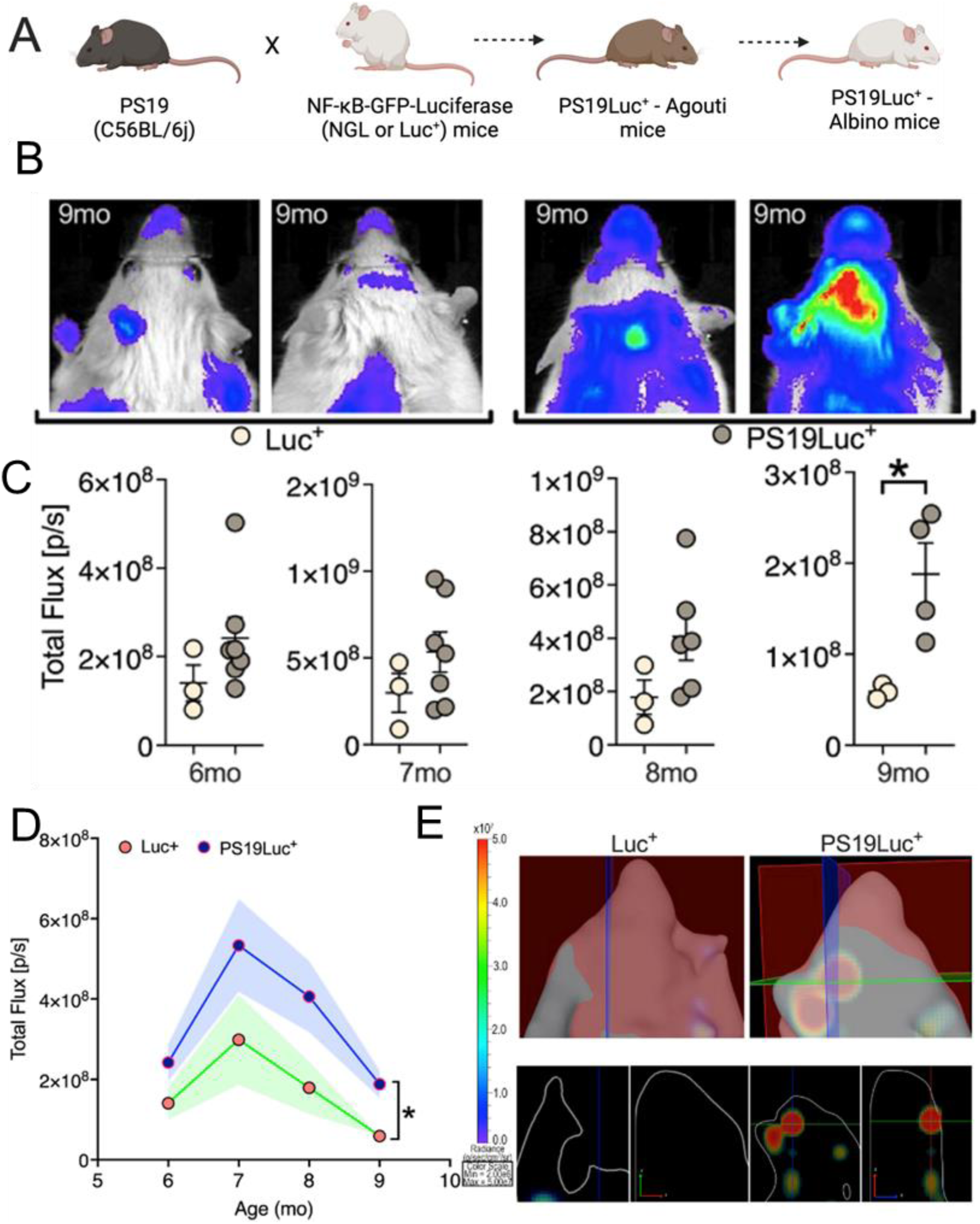
NF-κB activation in the head region is elevated as early as 6 months of age, but significantly higher at 9-month-old PS19Luc^+^ mice. **(A)** Schematic showing the breeding scheme to obtain Luc^+^ and PS19Luc^+^ mice in albino background for IVIS imaging. **(B)** Representative IVIS scans of the head region showing PS19Luc^+^ mice show an overall higher level of head-region bioluminescence signal indicating NF-κB activation as early as 6 months of age compared to Luc^+^ mice but was significantly higher at 9 months of age. **(C, D)** Individual mouse and mean+SEM of each group (D) of total flux (bioluminescence signal indicating NF-κB activation) in the head regions of Luc^+^ and PS19Luc^+^ mice at different ages (in months). **(E)** Representative 3D images showing that the bioluminescence signal in the head region of PS19Luc^+^ mice was localized to deeper (likely from the brain) tissue (right bottom panel). Data displayed as mean + SEM, Two-way ANOVA with Sidak’s multiple comparison test or unpaired t-test, *p<0.05, with n=3-7).

PS19Luc^+^ mice were subjected to longitudinal IVIS imaging at 6-, 7-, 8-, and 9-months of age to determine how age-related tau pathology in PS19 mice affects NF-κB activation. First head region field of view (FOV) analyses showed a robust bioluminescence signal in 9-month-old PS19Luc^+^ mice compared to age-matched littermate Luc^+^ control mice (Fig. 5B). Quantification of total flux showed that the head region FOV was elevated as early as 6-months of age, and statistically significant at 9-month-old PS19Luc^+^ mice (Fig. 5C-D). Bioluminescence signal from individual Luc^+^ and PS19Luc^+^ mice used in this experiment are shown in Supplemental Fig. 2A-B. Elevated bioluminescence signal in PS19Luc^+^ mice at 6- and 9-months of age, which are much higher than the age-matched Luc^+^ littermate control mice, is very evident. Next, we changed the FOV for the whole-body quantification at 7- and 9-month-old Luc^+^ and PS19Luc^+^ mice, which show significantly higher elevated whole-body bioluminescence in PS19Luc^+^ mice (Supplemental Fig. 2C-E). Finally, we performed 3D analysis to determine if the bioluminescence signal was superficial or coming from the deeper tissues. Spectrum 3D rendering showed that the signal was coming from the deeper tissues in the head region (likely from the brain) and whole-body (Fig. 5E, Supplemental Fig. 2F, and Supplemental Movie 1). Together, these results suggest both brain-specific and whole-body NF-κB activation in PS19Luc^+^ mice, which starts as early as 6-months and increases until 9-months. Due to the time needed to generate sufficient PS19Luc^+^ mice and to standardize IVIS imaging protocol, we could not perform any IVIS imaging on ice earlier than 6 months of age, which will be planned in future studies.

### NF-κB activation in the head region peaks at 10- and 12 months of age in hTauLuc^+^ mice

To validate our observations made in PS19Luc^+^ mice, we also crossed Luc^+^ mice with hTau mice (which expresses all six isoforms of human non-mutant tau protein under the endogenous human tau promoter and knockout for endogenous mouse tau)^74^ to generate albino hTauLuc^+^ mice (Fig. 6A). Unlike PS19 mice, the hTau mice show tau hyperphosphorylation starting at 3 months of age, Gallyas silver positive NFTs at about 9-12 months, and they live much longer than PS19 mice. We confirmed the presence of human tau in hTauLuc^+^ mice via genotyping (Supplemental Fig. 5D). We also confirmed that like 11-month-old hTau mice, 11-month-old hTauLuc^+^ mice show impairment in delayed-dependent memory in novel-object recognition test (Supplemental Fig. 1D). Because of slower onset and progression of tau pathology, we performed IVIS imaging in hTauLuc^+^ mice every month starting at 9-through 15-months of age. While 9- and 10-month-old hTauLuc^+^ mice did not show any increase in the bioluminescence in the head-region FOV, the signal peaked at 11- and 12-months of age when the NF-κB activity was significantly higher in hTauLuc^+^ mice (Fig. 6B-D). Spectrum 3D IVIS image analyses suggested signal origin from the deeper tissues (Fig. 6E, and Supplemental Movie 2). However, unlike PS19Luc^+^ mice, we did not observe a systemic elevation of bioluminescence signal when the FOV analysis was set for whole body in hTauLuc^+^ mice (Supplemental Fig. 3A-D). These results suggest longitudinal systemic and head-region NF-κB activation correlate with the severity of tau pathology in hTau and PS19 mice.

**Fig. 6.**
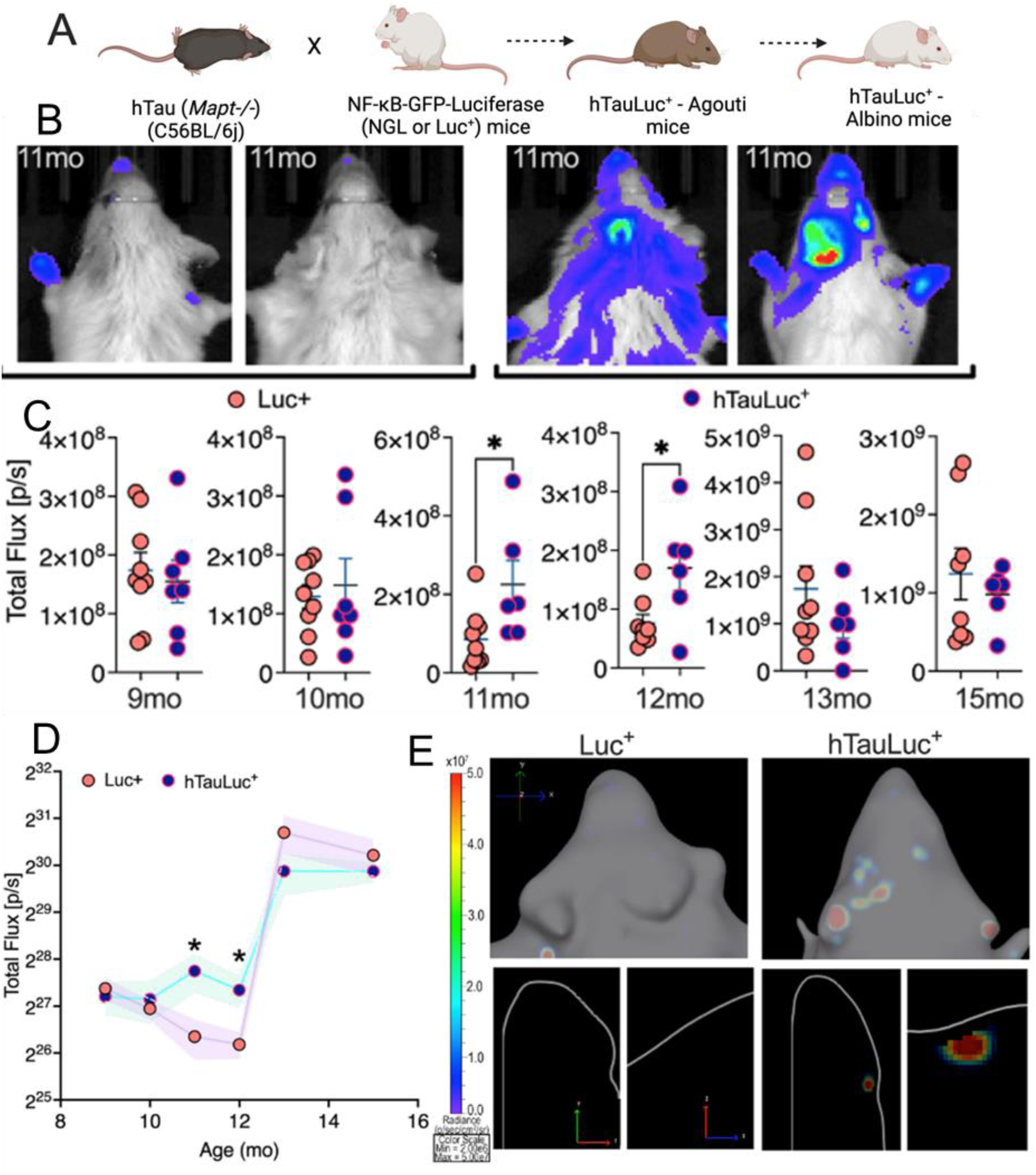
NF-κB activation in the head region peak at 10- and 12-months of age in hTauLuc^+^ mice. (A) Schematic showing the breeding scheme to obtain Luc^+^ and hTauLuc^+^ mice in albino background for IVIS imaging. **(B)** Representative IVIS scans of the head region showing hTauLuc^+^ mice show overall higher level of head-region bioluminescence signal indicating NF-κB activation at 11 months of age compared to Luc+ mice. **(C, D)** Individual mouse and mean+SEM of each group (D) of total flux (bioluminescence signal indicating NF-κB activation) in the head regions of Luc^+^ and hTauLuc^+^ mice at different ages (in months). **(E)** Representative 3D images showing that the bioluminescence signal in the head region of hTauLuc^+^ mice was localized to deeper (likely from the brain) tissue (right bottom panel). Data displayed as mean + SEM, Two- way ANOVA with Sidak’s multiple comparison test or unpaired t-test, *p<0.05, with n=7-9).

### Reducing pTau levels chemically (via Doxycycline) reduces NF-κB activation

In prior studies, we have demonstrated that lowering pathological tau levels in the rTg4510 doxycycline (DOX) regulatable tauopathy mouse model significantly decreases inflammasome markers (NLRP3 and ASC)^5^. To determine whether reducing the expression of human P301L tau in rTg4510 mice affects NF-κB activation, we maintained pregnant rTg4510 female mice on DOX diet until they deliver the pups. Once the pups are born, dams and litters were maintained on DOX diet until the litters were 2 months of age. Then, they were sacrificed to assess the levels of total human tau, phosphorylated IKKα/β and phospho-p65. Control rTg4510 mice were maintained on a regular diet, which led to the expression of human P301L tau. DOX-treated rTg4510 mice showed expected reduction Tau12^+^ human tau in the hippocampal lysates (Supplemental Fig. 5A-B). Notably, DOX-treated rTg4510 mice also showed a significant decrease in p-IKKα/β and p-p65 compared to rTg4510 mice treated with regular diet (Supplemental Figure 5A-B), suggesting the suppression of human mutant tau expression in rTg4510 mice reduces markers of NF-κB.

### Tau knockout mice show reduced NF-κB activation in the head region

In the second set of experiments, we crossed Luc^+^ mice to the tau knockout mice (*Mapt^-/-^*) to generate albino *Mapt^-/-^*Luc^+^ mice (Fig 7A). In several previous studies, we have reported that inducing acute systemic inflammation with LPS tau hyperphosphorylation^10^. The goal of this next experiment is not to test LPS-induced tau phosphorylation but to determine whether or not the endogenous mouse tau contributes to the LPS-induced systemic- and head-region (brain) inflammation. LPS (5 mg/kg b.w; single dose, i.p; for 3h) induced NF-κB activation (bioluminescence) was evident near the injection site in the peritoneal cavity and head-region of Luc*^+^*mice (Fig. 7B). Strikingly, bioluminescence signal was significantly reduced in both head-region and peritoneal area of *Mapt^-/-^*Luc^+^ mice when thresholded for the Luc^+^ mice bioluminescence (Fig. 7B). Note that these mice were imaged at the same time, side-by-side. Quantification of total flux in the head region showed significantly reduced NF-κB activity in LPS-injected *Mapt*^-/-^Luc^+^ mice compared to LPS-injected Luc^+^ mice (Fig. 7C). Notably, re-imaging the same mice at 24h after LPS injection did not show any difference in the bioluminescence or total flux in both groups of mice (Supplemental Fig. 4). Together, these results suggest that presence of tau and its pathological modification (hyperphosphorylation) upon LPS may likely contributes to the NF-κB activity in the brain.

**Fig. 7.**
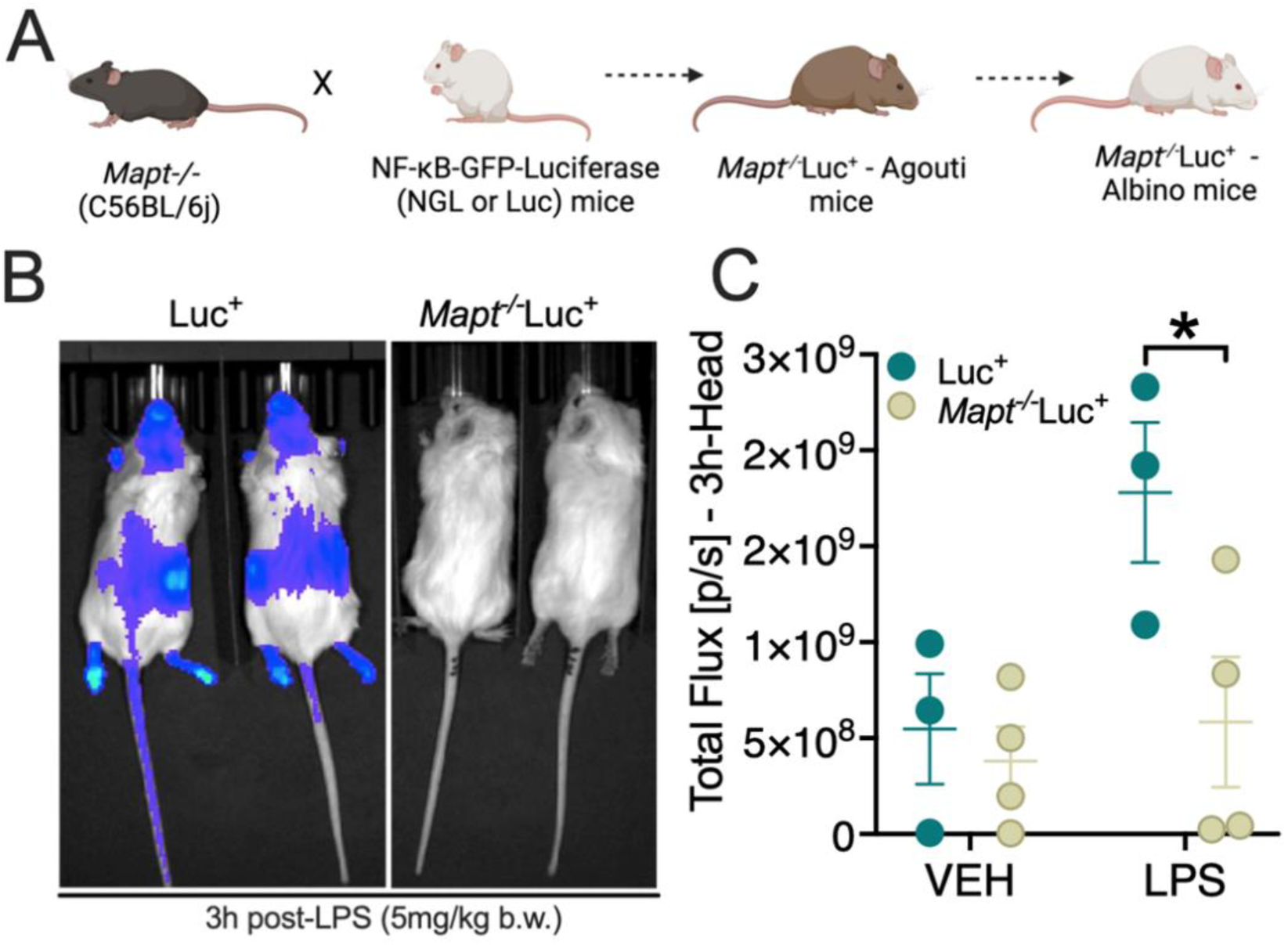
Tau knockout mice show reduced NF-κB activation in the head region. **(A)** Schematic showing the breeding scheme to obtain *Mapt*^-/-^/Luc^+^ and Luc^+^ *(Mapt*^+/+^/Luc^+^) mice in albino background for IVIS imaging. **(B)** Representative IVIS scans showing Luc^+^ mice show overall higher level of systemic and head-region bioluminescence signal at 3h after single dose of LPS (5 mg/kg b.w.; single dose; i.p) injection compared to LPS-injected *Mapt*^-/-^/Luc^+^ mice. **(C)** Quantification of total bioluminescence flux shows significantly reduced signal in the head region of *Mapt*^-/-^/Luc^+^ mice compared Luc^+^ mice. Data displayed as mean + SEM, unpaired t test or one-way ANOVA, *p<0.05, with n=3-4).

### Neutralization of pTau by Qβ-pT181-reduces NF-κB activation

While pharmacological (DOX-mediated) and genetic (*Mapt* knockout) based tau reduction showed blunted NF-κB activity, we next attempted to gain a proof-of-concept for future translatability of pathological tau reduction on NF-κB activity and overall neuroinflammatory responses. We vaccinated PS19Luc^+^ mice with our well-characterized Qβ virus-like particle (VLP)-based vaccines against phosphorylated T181 tau (Qβ-pT181). We have previously shown that Qβ-pT181 induces robust anti-pT181 antibody responses, safe, reduces soluble pTau and Sarkosyl insoluble/Gallyas silver positive NFTs/tau aggregates from the vaccinated rTg4510, PS19, and hTau mice^5,64,84^. We have also demonstrated that the anti-pT181 antibodies can enter the CNS, localize to neurons, show robust immunogenicity, and engage the target without adverse events in non-human primates (rhesus macaques)^84^. Qβ-pT181 vaccinated PS19Luc^+^ mice showed robust anti-pT181 antibody titers (Supplemental Fig. 5E) and significant reduction in the bioluminescence and total flux in the whole body and a >50% reduction in the head region compared to Qβ-control vaccinated PS19Luc^+^ mice (Fig. 8A-B). We confirmed the presence of human P301S transgene in both Qβ-control and Qβ-pT181 vaccinated PS19 mice via genotyping and PCR (Supplemental Fig. 5D). We validated that the reduction in NF-κB in Qβ-pT181 vaccinated PS19Luc^+^ mice by performing Western blot analyses, which showed significant decrease in AT108^+^ and Tau12^+^ pTau (and not difference in AT8^+^ and Tau5^+^ tau) (Supplemental Fig. 5), consistent with the reduced AT180 and Tau12 we reported recently in the original PS19 mice^84^. In a separate experiment, Qβ-pT181 vaccination also significantly reduced *Nfκb1*, *Nfκb2*, *Myd88*, *Nlrp3*, and *Pycard* (*ASC*) mRNA levels, with a modest reduction in the *Il1b* mRNA in original PS19 mice compared to Qβ-control vaccinated control PS19 mice (Supplemental Fig. 5C). Finally, post-necropsy assessment of p-p65 S536 and IκBα levels in Qβ-pT181 vaccinated PS19Luc^+^ mice hippocampi showed no changes in the p-p65 S536 levels (Fig. 8C-D). However, total IκBα levels significantly elevated in Qβ-pT181 vaccinated PS19Luc^+^ mice compared to Qβ-control vaccinated PS19Luc^+^ mice. These results suggest neutralizing pTau with anti-pT181 antibodies via Qβ-pT181 vaccination reduces NF-κB activity in PS19 mice.

**Fig. 8.**
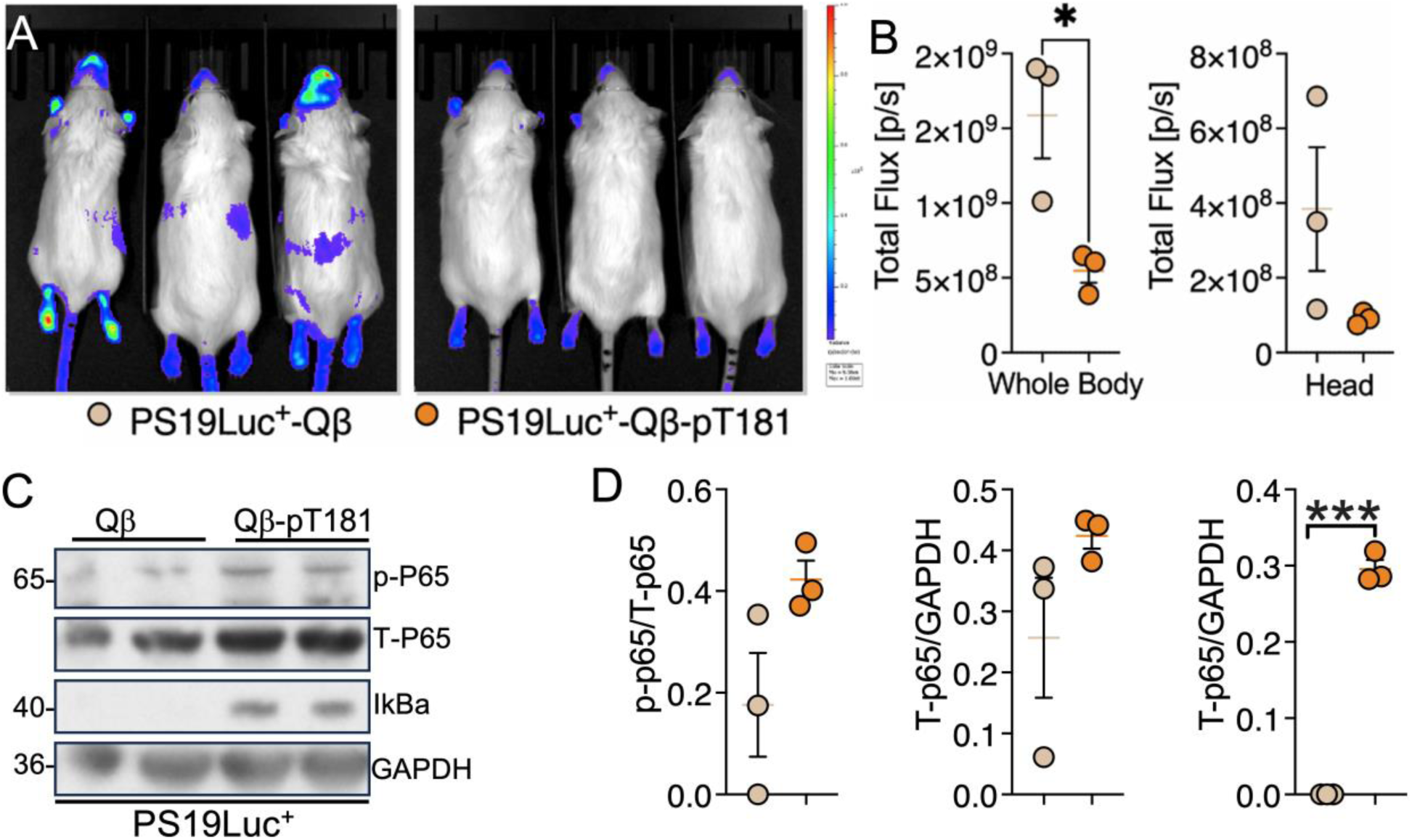
Neutralization of pTau by pT181-Qβ reduces NF-κB activation. **(A)** Representative IVIS scans show an overall higher level of bioluminescence signal indicating NF-κB activation in albino PS19Luc^+^ with Qβ control vaccinated mice compared to PS19Luc^+^ with Qβ-pT181 vaccinated mice. **(B)** Total bioluminescence flux (NF-κB activation) is markedly reduced in the whole body of Qβ-pT181 vaccinated PS19Luc^+^ mice compared to Qβ control injected PS19Luc^+^ mice. Note that the head region shows a four-fold reduction in the bioluminescence signal. **(C, D)** Western blot and quantification showed significantly increased IκBα levels, which indicates decreased NF-κB activation in Qβ-pT181 vaccinated PS19Luc^+^ mice compared to Qβ control injected PS19Luc^+^ control mice. Data displayed as mean + SEM, unpaired t-test, *p<0.05, with n=3.

## Discussion

We provide a comprehensive analysis of the pro-inflammatory roles of pathological tau via activation of NF-κB. NF-κB is one of the master regulators of innate immune activation and is known to contribute to chronic neuroinflammation in AD and other tauopathy. Tau oligomers drive NF-κB activation more than PHFs, and via chronically promoting the degradation of IκBα. Apparently, AD-TO induced IκBα degradation appears to be independent of major proteostatic process of autophagy, proteasomes and calpain. Rather, the IκBα secreted into the extracellular space upon AD-TO stimulation of C20 microglia. We provide evidence for pathological tau-induced longitudinal NF-κB activation in the head-region and whole body of PS19Luc^+^ and hTauLuc^+^ mice corresponding to the progressive nature of tau pathology in these mice. Finally, we provide evidence that suppression of pTau via pharmacologically (DOX in rTg4510 mice), genetically (via tau knockout in Mapt^-/-^ Luc^+^ mice with LPS) and via antibody-mediated neutralization (Qβ-pT181 vaccinated PS19Luc^+^ mice) reduces NF-κB activation.

NF-κB activation is a rapid cellular defense response to thwart invading pathogens or other ‘tissue damaging elements’ by immune cells^62^. However, chronic NF-κB activation can result in inflammation because NF-κB is a master regulator of numerous pro-inflammatory cytokines, chemokines and immune factors ^63^. Our findings corroborate with other reports where microglia-specific NF-κB activation has been shown to promote microglia-dependent tau seeding and spreading in PS19 mice^34^, and blocking microglial NF-κB prevents tau seeding/spreading and improves memory. Likewise, Aβ40 peptide was shown to induce dimerization of p50/p65 and can lead to NF-κB activation^64^. Together, these studies highlight the role of NF-κB activation in triggering chronic neuroinflammation in AD and related tauopathy.

Since the brains of non-amyloid tauopathy, where pathological tau accumulation is typically intra-neuronal, also show evidence of significant inflammation 3-5, it is pivotal to determine how neuronal pathological tau chronically elicits the innate immune response is mediated by NF-κB. Our previous findings7,8 and results from this study provide compelling evidence that pathological tau exacerbates neuroinflammation by activating the NF-κB and IL-1β axis^5,10,64^. An earlier study has shown that NF-κB can induce the expression of SET gene isoform 1, which is upregulated in AD brains and contains a functional κB sequence in its promoter region and known to inhibit protein phosphatase 2A (PP2A) that dephosphorylates tau ^65^. Other studies have demonstrated that Aβ can induce calcineurin activation, leading to NF-κB induced activation of mGluR5, which accumulates near Aβ plaques^66^. Therefore, NF-κB activation has repeatedly been implicated both in AD and other non-AD tauopathies where it can drive chronic neuroinflammation via various mechanisms.

Our cell culture results suggest that AD-TO induced IκBα degradation is independent of major proteostatic pathways but promotes the secretion into the extracellular space via most likely secretory autophagy, significantly when proteasome, autophagy and calpain are inhibited. The exact mechanisms involved in the extracellular secretion of IκBα and its role in the extracellular space are unclear. While the tri-inhibition of major proteotoxic pathways seems non-physiological, one or more of these pathways have been reported to be compromised in the brains of AD and other tauopathies^26,27,67^.

NF-κB activation in PS19Luc^+^ and hTauLuc^+^ mice aligns with the appearance and progression as well as severity of tau pathology, the limitation of our study is that we did not perform analyses any earlier than 6 months of age in case of PS19Luc^+^ mice and 9 months of age in hTauLuc^+^ mice. Nonetheless, IVIS imaging results correlate with the post-necropsy Western blot analyses, and data from removal of pathological tau by DOX, *Mapt* deficiency and anti-pT181 antibody-based neutralization. We know that DOX can itself serve as an anti-inflammatory besides an antibiotic. However, consistency of reduced NF-κB activity in tau knockouts in LPS model and Qβ-pT181 vaccinated PS19Luc^+^ mice suggests that the DOX-study finding indeed is accurate. Likewise, we are not uncertain why the IVIS imaging quantification shows a peak in the NF-κB activity in both hTauLuc^+^ and PS19Luc^+^ mice. Still, their either reduces at later age (in PS19Luc^+^ mice), and elevated in hTauLuc^+^ mice. We are unclear why NF-κB decreases and increases with age in PS19 vs. hTau mice. Some activated microglia may undergo pyroptosis, leading to reduced NF-κB in PS19 mice, and hTau mice may have different NF-κB regulation than PS19. This possibility needs to be tested in future studies.

In LPS-treated *Mapt*^-/-^Luc^+^ mice, the 3h time point shows a significant reduction in NF-κB activity, which returns to baseline by the 24h time point. We speculate that the 24h time point reduces either the acute activation of NF-κB at the 3h time point, unless the *Mapt*^-/-^Luc^+^ mice show LPS-induced NF-κB activation at a much slower rate due to loss of tau than LPS-injected Luc^+^ mice. Qβ-pT181 vaccinated PS19Luc^+^ mice showed reduction in AT180 positive pathological tau, similar to the Qβ-pT181 vaccinated PS19 mice that we recently published^84^. It is unclear if only AT180 positive, and not AT8 positive, tau is specifically sensitive to anti-pT181 antibody mediated neutralization. Upon reduction of this pathological tau species is sufficient to correlate with reduced NF-κB activation. One limitation of our study is that we have not investigated other pathological forms of tau (oligomers, PHFs, and truncated tau), which will be examined in future studies. Finally, we observed that Qβ-pT181 vaccinated PS19Luc^+^ mice specifically increased IκBα levels without significantly reducing p-p65 S536 levels. Because reduction in IκBα correlated with the IVIS imaging data on reduced NF-κB activation, we conclude that IκBα levels are a good surrogate marker of NF-κB activation rather than p-p65 S536 levels, specifically in PS19Luc+ mice.

In conclusion, our results suggest that pathological tau has a pro-NF-κB activation role and thus contributes to neuroinflammation in AD and related tauopathy. Upon further validation of the time points of peak NF-κB activation reported here in independent studies, prophylactic vaccination with Qβ-pT181 and neutralization of pTau may prevent pTau-induced NF-κB activation and thus provide a means to reduce neuroinflammation in AD and other tauopathy.

## Materials and Methods

### Pathological tau isolation from autopsy of human tauopathy brains and characterization

Tau monomer and oligomers were isolated from autopsy of human frontotemporal lobar degeneration (FTLD)-associated brain were a kind gift from Dr. Rakez Kayed, UTMB Galveston, Texas. The structural morphology of tau monomer and oligomers was characterized using a super-resolution direct stochastic optical reconstruction microscopy (dSTORM). Transmission electron microscopy (TEM) was used to characterize neurofibrillary tangles containing straight and paired helical filaments at nanoscale resolution.

### Purification of Sarkosyl insoluble paired helical filaments (PHFs) from human AD brain

The Greenberg and Davies (1990)^68^ protocol was modified to isolate paired helical filaments (PHFs), which was also previously described by us^5^. Human AD (Braak stage VI) brains were homogenized in buffer H, sonicated, and centrifuged at 22,000 g for 30 min at 4°C. The supernatant was adjusted to 1% N-Laurylsarcosine and 1% β-Mercaptoethanol, incubated at 37°C for 2 h with agitation, then centrifuged at 150,000 g for 35 min. The Sarkosyl-insoluble pellet was washed and resuspended in Buffer H with 1% CHAPS and 1% β-Mercaptoethanol. After filtration and centrifugation, the PHF pellet was resuspended in Buffer H with β-Mercaptoethanol and purified via sucrose gradient centrifugation (35,000 rpm, 2 h, 4°C). PHFs were collected from the 35% layer and interface. Further purification was done using chloroform/methanol precipitation to remove sucrose. The protein sample was mixed with methanol and chloroform, vortexed, and centrifuged at 10,000 g for 5 min to isolate PHFs at the phase boundary. Additional methanol washes and centrifugation followed, and the final PHFs pellet was air-dried and resuspended in cell culture media. All steps were performed at room temperature.

### Direct stochastic optical reconstruction microscopy (dSTORM)

dSTORM super-resolution microscopy for tau monomer and oligomers was done as previously reported^69,70^. First, the tau monomer and tau oligomers (AD-TO) were labelled with Alexa Fluor 647 NHS Ester by mixing them at 5:1 ratio in PBS (pH, adjusted by 1M NaHCO3). The reaction mixture was incubated overnight at 4°C in the dark. Fluorescent labelled tau monomer or AD-TO were further purified by using a Zeba spin desalting column 7-kDa MCAO, (ThermoFisher # 89882). An average fluorophore: protein ratio was maintained at 2:1 as determined by the absorption spectroscopy. dSTORM imaging was done at an exciting intensity of ∼1 kW/cm2. The fluorescence filter setup consisted of a dichroic mirror (650 nm; Semrock, Rochester, NY) and an emission filter (692/40; Semrock). Additional details of super resolution image reconstruction and mathematical modeling for single emitter testing in MATALB are reported in a previous published study^72,73,74^ from a collaborator team who helped with the dSTORM imaging at the University of New Mexico.

### Transmission electron microscopy (TEM)

Purified PHFs from human AD brains were placed on a carbon coated transmission electron microscopy (TEM) grids as we reported previously^5^, and left on the grid for 5 min, excess solution removed by Whatman filter paper. Grids were stained with 2% uranyl acetate for 2 min before imaging the samples in a Hitachi H7500 TEM equipped with an Advanced Microscopy Sciences XR60 camera.

### Cell culture

Bone marrow macrophages (BMMs) 10^5^ cells/well from NF-κB-GFP-Luciferase (NGL) mice were grown and treated as we previously described^5^. Briefly, BMMs were prepared by flushing the femurs and tibia of 3-month-old NGL mice with a syringe and 24 G needle containing differentiation media, which include macrophage-colony stimulating factor (M-CSF) as previously described [10]. The cells were treated with fresh M-CSF media every 2 to 3 days. After this, the cells (BV2s and BMMs) were treated with LPS (from E coli 055: B5, Sigma # L2880-25MG, Lot # 102M4017V; 200ng/ml for 12 h), tau monomer, tau oligomers (AD-TO), PHFs treatment (at 2 µg/ml for 12h) and IVIS imaging. The BV2 microglial cell line was a gift from Dr. Gary Landreth. Human C20 microglia (a kind gift from Dr. David Alvarez-Carbonell)^71^ and mouse BV2 microglia cell lines were cultured in DMEM (ThermoFisher # 11965092) and supplemented with 10% heat-inactivated fetal bovine serum (FBS) (ThermoFisher # 16140071) and 1X concentration of penicillin-streptomycin (10,000 U/mL, stock) (ThermoFisher # 15140122). All the cell cultures were maintained at 37° C with 5% CO2 and moisture.

### NF-κB-GFP-Luciferase Assay

In a cell culture setting, the NF-κB-GFP-Luciferase Assay was performed as we previously reported for another luciferase assay^72^. Briefly, for firefly luciferase activities, D-Luciferin, potassium salt (ThermoFisher # L2916) was reconstituted in water and was added (1:100) to each well of BMM cells treated with tau monomers or AD-TO. The 24-well plate was imaged in the IVIS Lumina Series II with system software three to four minutes after adding the substrate. (n = 1 refers to an entire 24-well plate, and six wells individually calculated per control). The plate scans were thresholded to normalize to control levels and the output bioluminescence was expressed as relative light units.

### Proteostasis inhibitors treatment and sample preparation for western blotting

Microglia C20 cells (0.1 x 10^6^) were seeded in a 12-well plate. Next day, C20 cells were treated with AD-TO (1mg/ml) to stimulate NF-κB activity chronically for 48h. Protein translation was inhibited at 43h with 50µM of cycloheximide (CHX) 1h prior adding the proteostasis inhibitors at 44h, including 10µM MG132 (26S proteasome), 100nM Bafilomycin A1 (Baf A1, autophagy), 100µM PD 150606 (calpain 1/2) either individually or all three inhibitors together for 4h. Similarly, 20S proteasomal activity was inhibited with 10 µM lactacystin to identify IκBα as a substrate of 20S proteasome. All the inhibitors were reconstituted with DMSO to prepare stock solutions and stored at -20° C until further use. After treatment, cells were washed with 1X PBS, three times and lysed with ice cold RIPA buffer (ThermoFisher # 89901) with 1% PMSF (Sigma # 93482-50ML-F), 1X concentration of protease (Sigma # P8340-1ML) and phosphatase (Sigma # P2850-1ML) inhibitor cocktails and incubated on a rocker for 30min at 4° C. Then the cell lysates were centrifuged at 14,000g for 10min at 4° C. The resulting supernatant was stored at -80° C or heat-denatured the lysates with 1X LDS-sample buffer (ThermoFisher # NP0007) at 95° C for 10min. Samples were centrifuged to collect the supernatant at 14,000g for 5min at 4° C prior loading the denatured samples onto NuPAGE Bis-Tris protein gels for protein separation (ThermoFisher # NP0322BOX).

### Lipopolysaccharide (LPS) administration

A single dose of LPS (from E coli 055: B5, Sigma # L2880-25MG, Lot # 102M4017V) was administered to two-month-old Mapt^-/-^ and Mapt^-/-^Luc^+^ mice (in albino background) mice as previously described ^73^ [11]. Briefly, Vehicle (Veh, Hank’s Balanced Saline Solution) or LPS was administered at 5 mg/kg body weight (intraperitoneal; single dose). After 3h, the mice underwent IVIS imaging of the whole body and head regions (see below). Soon after IVIS imaging, the mice were sacrificed, and the left hemisphere hippocampi were processed for biochemical analysis.

### Tissue Preparation for Biochemical Analysis

Mice were anesthetized and transcardially perfused with 0.125 M phosphate buffer (PB). After perfusion, the brains were extracted. The left hemisphere was immersion fixed in 4% paraformaldehyde in PB (4% PFA/PB), while the right hemisphere was micro-dissected into the cortex and hippocampus. Wet weights were recorded, and tissues were snap-frozen in liquid nitrogen for biochemical analysis. The remaining right hemispheres were weighed and snap-frozen for mRNA extraction. Hippocampi from different mouse strains were used for bio-chemical analysis, including qRT-PCR and Western blot. Immunohistochemical analysis was performed on 30μm thick sagittal brain sections.

### RT-qPCR

RNA from cells and mouse brains was extracted using the TriZOL reagent described by the manufacturer (ThermoFisher). Total RNA (20 ng/µL) was converted to cDNA using the High-Capacity cDNA Reverse Transcription kit (ThermoFisher) and amplified using specific TaqMan assays (catalog # 4331182; ThermoFisher) (additional details in Supplemental Table S1). GAPDH (ThermoFisher # 4352339) and 18s rRNA (catalog # 4319413E, ThermoFisher) were used as a housekeeping gene for normalization. qRT-PCR assays were run on the StepOnePlus® Real-Time PCR System (ThermoFisher) and the statistical analyses were performed using GraphPad Prism. The PCR primer pairs utilized are indicated in the Supplemental Table S1.

### Western blotting

Cell or tissue lysates were diluted with a final concentration of 1x LDS/RA buffer and sonicated for 30 seconds, then heat denatured at 95°C for 10 minutes. Samples were centrifuged to collect the supernatant at 14,000g for 5min at 4° C prior loading the denatured samples onto NuPAGE Bis-Tris protein gels for protein separation (ThermoFisher # NP0322BOX). Separated proteins gel was blotted onto a PVDF Transfer Membranes, 0.2μm (ThermoFisher # 88520) in 1X Tris-Glycine transfer buffer containing 20% methanol for overnight at 4° C. Membrane was blocked with either 5% nonfat milk powder or 3% BSA in 1X PBS for 1h at room temperature depending on the antibody datasheet from the vendors. The dilutions of primary antibodies utilized are indicated in the Supplemental Table S1.

### Animals

The University of New Mexico (UNM) Institutional Animal Care and Use Committee (IACUC) ap-proved all animal procedures described in the study under the IACUC protocols #s: 22-201247-B-HSC (Breeding); 21-201179-HSC (Experimental)). All mice were housed in a specific pathogen-free (SPF) facility within the Animal Research Facility (ARF) at UNM, in a 12h light/dark cycle with ad libitum access to food (Envigo’s 2920 diet) and water. Eighty-five square inch ventilated micro isolator cages were used to house the mice.

They were supplemented with sterilized and autoclaved TEK fresh standard crinkle bedding; environmental enrichment included tissue paper, wooden twigs, and an elevated penthouse insert. Each cage housed about 2-5 mice and grouped them by sex. Mice assigned for experiments were healthy with average weight, and had no history of medical conditions, including rectal prolapse, malocclusion, and dermatitis. The following mouse lines were utilized in the present studies: 1. PS19 mice^74^ – The Jackson Laboratory Strain # 024841, 2. hTau mice^75,76,77^, 3. NF-κB-GFP-Luciferase^78^ (NGL or Luc; The Jackson Laboratory Strain # 027529) mice, 4. Tau knockout (*Mapt^-/-^*) mice^81,82^, 5. rTg4510 mice^79^. Each of these mice strains was crossed to generate PS19Luc^+^ and hTauLuc^+^ mice until we obtained these NF-κB reporter mice with human tau transgenic background with white fur. The presence of human tau transgene and NF-κB reporter gene were confirmed by genotyping using the suggested primers from The Jackson Laboratory.

### Longitudinal IVIS imaging *in vivo*

NGL or Luc^+^, PS19Luc^+^ and hTauLuc^+^ mice were injected (i.p) with D-Luciferin (150 mg/kg body weight; Revvity; Cat # 122799) for 15 min prior to anesthesia by inhalation of isoflurane. Then the animals were imaged longitudinally at 9-, 10-, 11-, 12-, 13- and 15-months of age for hTauLuc^+^, and 6-, 7-,8-, and 9-months of age for PS19Luc^+^ along with age-matched control littermate Luc+ mice in the Perkin Elmer Spectrum Optical In Vivo Imaging System (IVIS). Scans were obtained for the field of view (FOV) of the whole body and the head region. In a few representative mice were also subjected to 3D imaging to assess the origin of signal from deeper tissues. For *Mapt*^-/-^ and *Mapt*^-/-^Luc^+^ mice, the IVIS imaging was done similarly, but after 3 h of LPS injection. The mice are placed on the heated platform during the imaging length to maintain the body temperature. The IVIS resolution was close to 20 µm (3.9 cm FOV). Total flux (normalized across group of mice from different groups imaged simultaneously) was calculated and expressed as mean + SEM of photons/second (p/s).

### DOX treatment

Doxycycline (DOX)-mediated regulatable rTg4510 (expressing human tau with P301L mutation) mice (Santacruz et al., 2005) were obtained from the Jackson Laboratory. Pregnant female mice were maintained in DOX diet (TD.05125, 2.1 g DOX/day ad libitum in chow, Tekland/Envigo, USA) until they gave birth to litters. Dams were continued on DOX diet until the litters were weaned. Then the litters were maintained in DOX diet until they are 2-months-age before sacrificing them for tissue analyses for Western blotting.

### Vaccination of Qβ-pT18 to hTau and PS19 mice

Qß-virus-like particles (Qß-VLPs or Qß) were produced in Escherichia coli (E. coli) and conjugated to ^175^TPPAPKp**T**PPSSGE<u>GGC</u>^190^ (peptide encompassing threonine 181 in human tau) as previously described^64^. Two-month-old PS19Luc^+^ mice received unconjugated control vaccination Qß-VLP or Qß-VLP conjugated to a pTau peptide (Qß-pT181) at 5µg/injection into the rear hind paw. Mice were injected bi-weekly, two doses, and allowed to age for two months after the last injection. The mice were sacrificed, and the hippocampi were processed to assess the levels of AT8+ and AT180+ pTau, phospho-P65 Ser536, total P65, total IκBα, human tau (Tau12) and GAPDH by western blots.

### Magnetic resonance imaging (MRI) scan of mice

Mice were imaged using a 7.4 Tesla Biospec MRI scanner (Bruker Biospin; Billerica) equipped with a single-tuned surface coil for the mouse brain. Briefly, mice were anesthetized with isoflurane gas (induction dosage 2-3%; maintenance dose 1.5-2%) delivered in a mixture of O2:NO2 gases in the ratio of 2:1 during the recording period (approximately 1-3 hrs). T2-weighted (T2w) images to assess cortex (CX) and hippocampal volumes were obtained. To evaluate volumetric changes, T2w images were minimally processed, and using the spline tool on VivoQuantTM (inviCRO, LLC, Boston, MA) and the Allen Brain Atlas as a reference (www.allenbrainatlas.org), regions of interest (ROI) were hand-drawn around the CX, HP and CC on every MR brain slice (n = 18 slices, per mouse). Additional imaging processing and analysis was performed using Para-vision (ver. 5.1, Bruker, Germany), the Bruker plugin available on FIJI Image J (NIH, USA), and VivoQuant software.

### Novel object recognition test

Novel object recognition task behavioral test was carried out as previously published^64^. Briefly, on the first day of this 3-day paradigm animals were acclimated to an open 75-cm2 arena for 5 minutes. On the second day, they were exposed to two identical objects (2 glass jars) for 5 minutes. On the test day, they are exposed to one of the previous familiar objects (glass jar) and a novel object (plastic water bottle). The ratio of novel (on the bottom right side of the maze) or familiar (top right of the maze) object encounters to total object encounters was recorded as a measurement of recognition memory.

### Statistics

Comparisons between the two groups were done via unpaired t test; comparisons between multiple treatment groups were done via one-way or two-way analysis of variance (ANOVA) with indicated multiple comparisons post-hoc tests. All statistical analyses were performed using GraphPad Prism® (Version 6.0).

## Supporting information

Tangavelou_etal_Supplemental Information

Movie1_Luc

Movie1_PS19Luc

Movie2_hTauLuc

Movie2_Luc

## Author Contributions

K.B., and B.C – designed the study. K.T., S.J., S.D., J.P.H., K.S., V.B., Z.V., M.M., and J.P – performed the experiments and analyzed the data. K.T – responsible for designing IκBα degradation/secretion experiments and data analyses. J.P.H., J.P., and B.C – responsible for pT181 Qβ synthesis and vaccination. K.T., and K.B - wrote the manuscript with input from all authors. Authors have read and agreed to publish the manuscript.

## Acknowledgments

We thank Ms. Irina Lagutina for performing all IVIS imaging and quantification at the Animal Model Core facility at the University of New Mexico. We also thank Dr. Keith Lidke and Dr. Aaron Neuman at the University of New Mexico for their help with the Direct stochastic optical reconstruction microscopy (dSTORM) images of tau monomers and oligomers. We also thank Dr. Rakez Kayed from the University of Texas Medical Branch at Galveston for providing the AD-TO purified from human AD brains.

