## Supplementary material for "Pathological tau activates inflammatory nuclear factor-kappa B (NF-κB) and pT181-Qβ vaccine attenuates NF-κB in PS19 tauopathy mice": Tangavelou_etal_Supplemental Information

*Article*

### Supplementary Figures

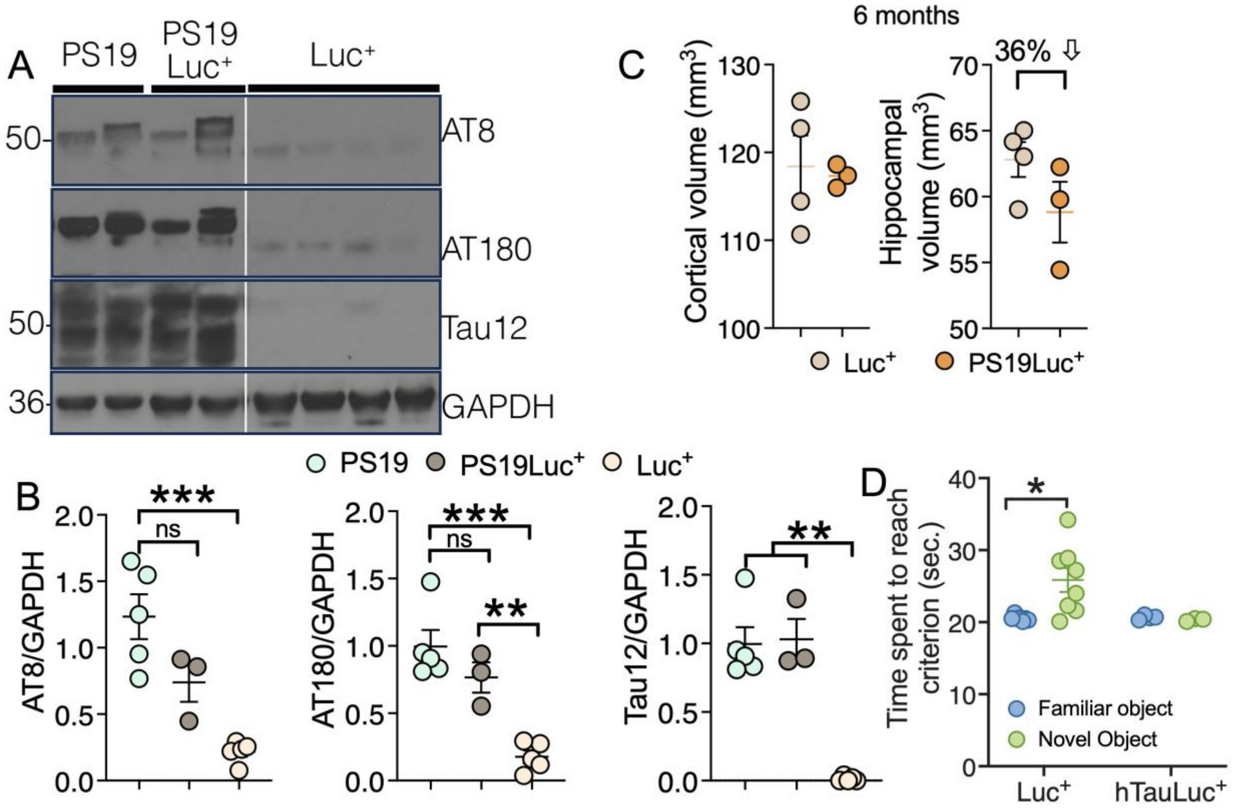

**Supplemental Fig 1: PS19Luc⁺ mice show elevated tau pathology similar to original PS19 mice and hTauLuc⁺ mice show impaired recognition memory at 11 months of age. (A, B)** Western blot analyses and quantifications (B) of albino 9-month-old PS19Luc⁺ mice in mixed genetic background shows elevated levels of human tau (Tau12) phosphorylated tau (pT231 or AT180⁺; pT202/pT205 or AT8⁺) similar to original 9-month-old PS19 mice in C57Bl/6j background. Nine-month-old Luc⁺ mice (littermate) showing basal phosphorylated or total tau levels are shown for comparison. **(C)** T2 volumetric MRI analyses show a 36% reduction in the hippocampal, but no difference in the cortical volume, in PS19Luc⁺ mice compared to age-matched Luc⁺ mice. **(D)** Novel-object recognition test shows impaired recognition memory in 11-month-old PS19Luc⁺ mice Data displayed as mean + SEM, unpaired t test or One-way ANOVA with Tukey's multiple comparison test and Two-way ANOVA with Sidak's multiple comparison test or unpaired t test, *p<0.05, with n=4-8).

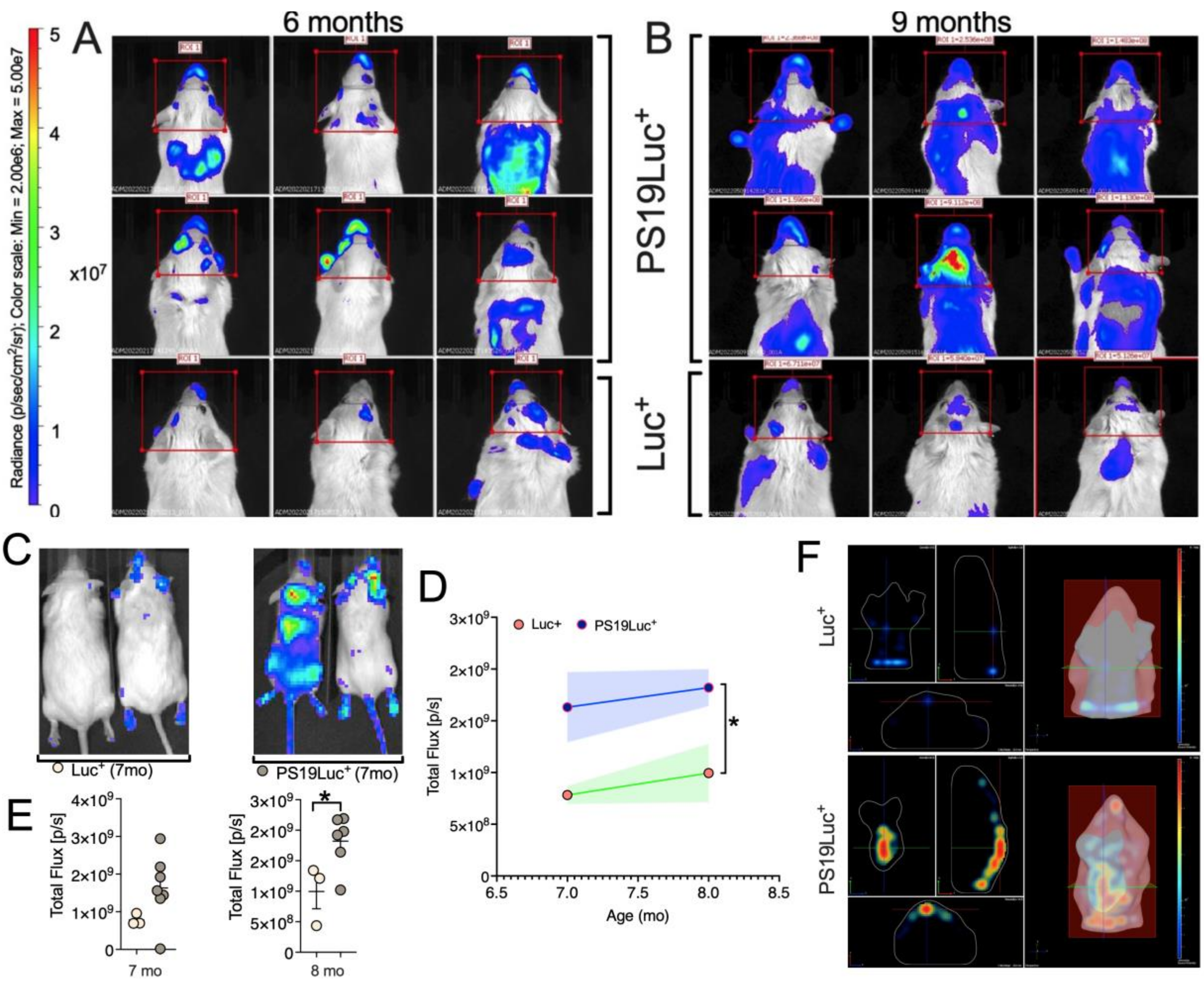

**Supplemental Fig 2: NF-κB activation in the whole body at 9-15-months of age in PS19Luc$^+$ mice. (A, B)** Representative IVIS scans of individual Luc$^+$ and PS19Luc$^+$ mouse in each group at each time point showing the head region bioluminescence signal indicating NF-κB activation as early as 6 months of age compared to Luc$^+$ mice but was significantly higher at 9 months of age. **(C-E)** Representative IVIS scans of the whole-body showing significant increase in the whole-body bioluminescence signal (NF-κB activation = Total Flux) in the from 7- months until 8 months in PS19Luc$^+$ mice compared to age-matched Luc$^+$ littermates. **(F)** Representative 3D images show the bioluminescence signal in the whole body (till torso) of Luc$^+$ and PS19Luc$^+$ mice showing general systemic NF-κB activation. Data displayed as mean + SEM, Two-way ANOVA with Sidak's multiple comparison test or unpaired t-test, *$p<0.05$, with n=7-9).

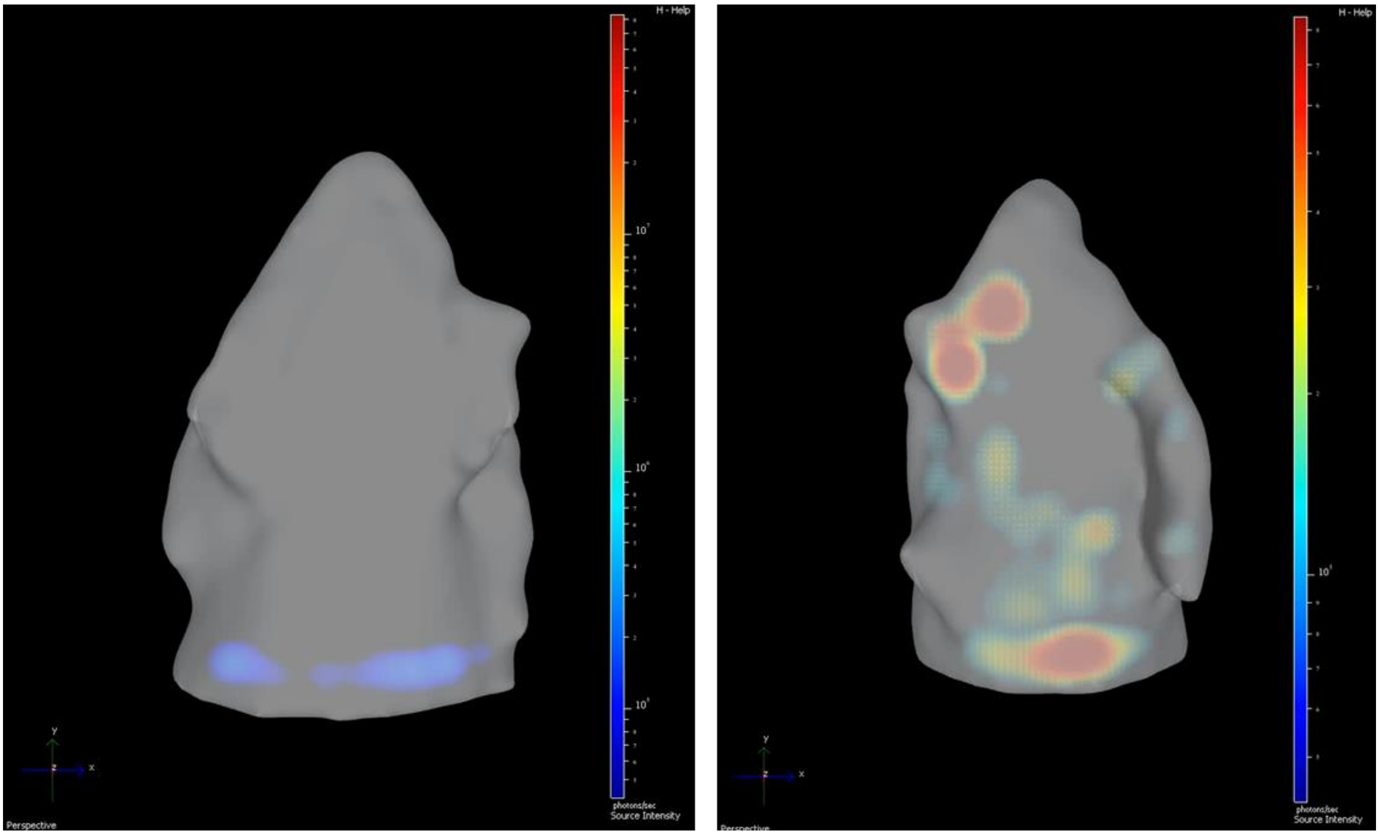

Luc+ PS19Luc+

**Supplemental Movie 1: Three-dimensional rendering of 9-month-old Luc+ and PS19Luc+ mice show localization of NF-κB signal in the brain region.**

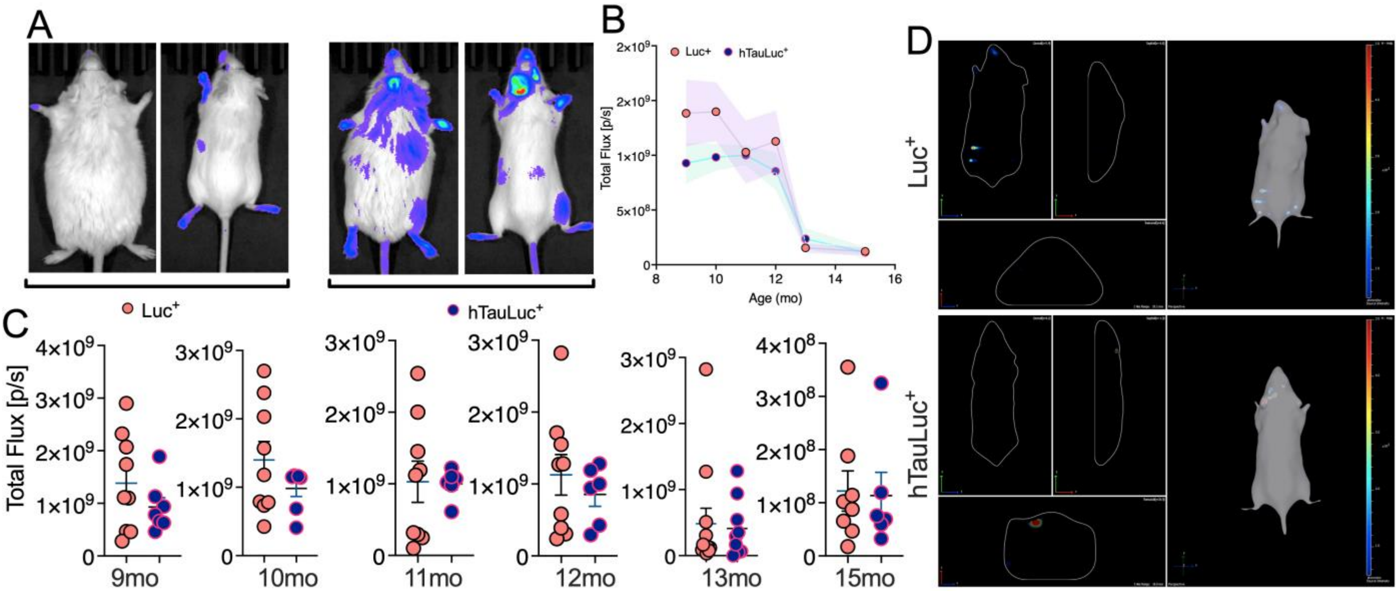

**Supplemental Fig 3: NF-κB activation in the whole body at 9-15-months of age in hTauLuc+ mice. (A-C)** Representative IVIS scans of the whole-body showing no significant difference in the bioluminescence signal (NF-κB activation = Total Flux) in the Luc+ and hTauLuc+ mice from 9- to 11 months of age. **(D)** Representative 3D images show the bioluminescence signal in the whole body of Luc+ and hTauLuc+ mice. Data displayed as mean + SEM, Two-way ANOVA with Sidak's multiple comparison test or unpaired t-test, *$p<0.05$, with n=7-9).

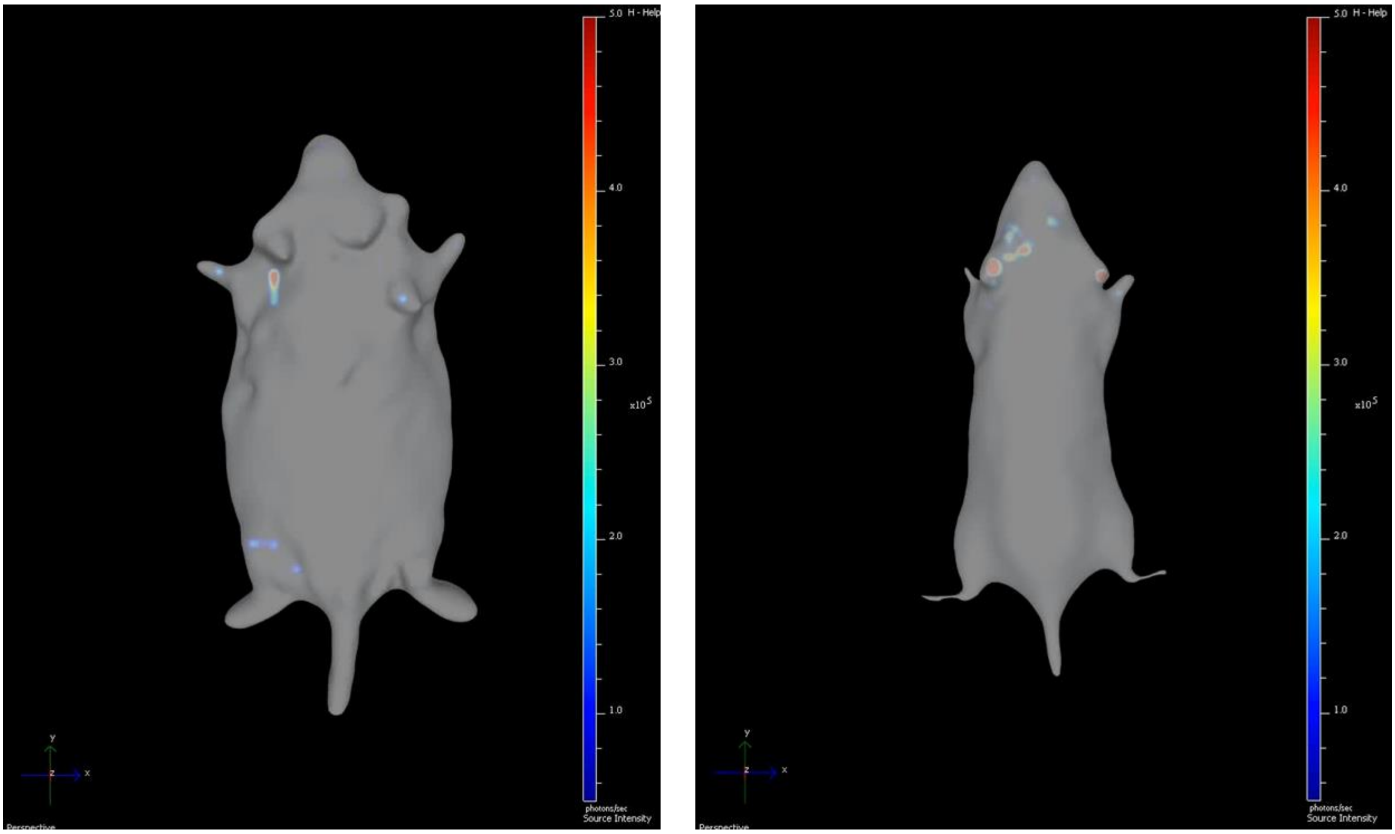

**Luc$^+$** **hTauLuc$^+$**

**Supplemental Movie 2: Three-dimensional rendering of 11-month-old Luc$^+$ and hTauLuc$^+$ mice show localization of NF-κB signal in the brain region.**

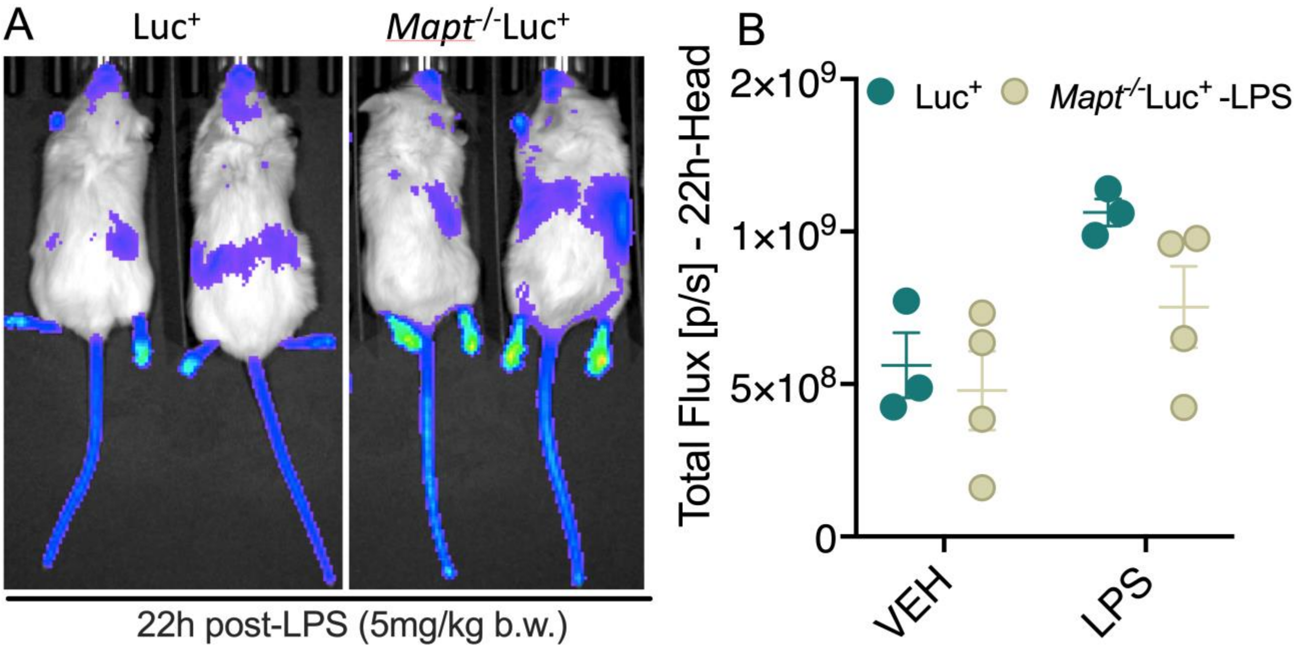

**Supplemental Fig 4: Tau knockout mice no change in the NF-κB activation by 22h post LPS injection.** **(A)** Representative IVIS scans showing Luc$^{+}$ and *Mapt*$^{-/-}$/Luc$^{+}$ mice show no difference in the systemic and head-region bioluminescence signal at 22 h after single dose of LPS (5 mg/kg b.w.; single dose; i.p) injection. **(B)** Quantification of total bioluminescence flux shows no difference in the signal in the head region of Luc$^{+}$ and *Mapt*$^{-/-}$/Luc$^{+}$ and mice. Data displayed as mean + SEM, n=3-4).

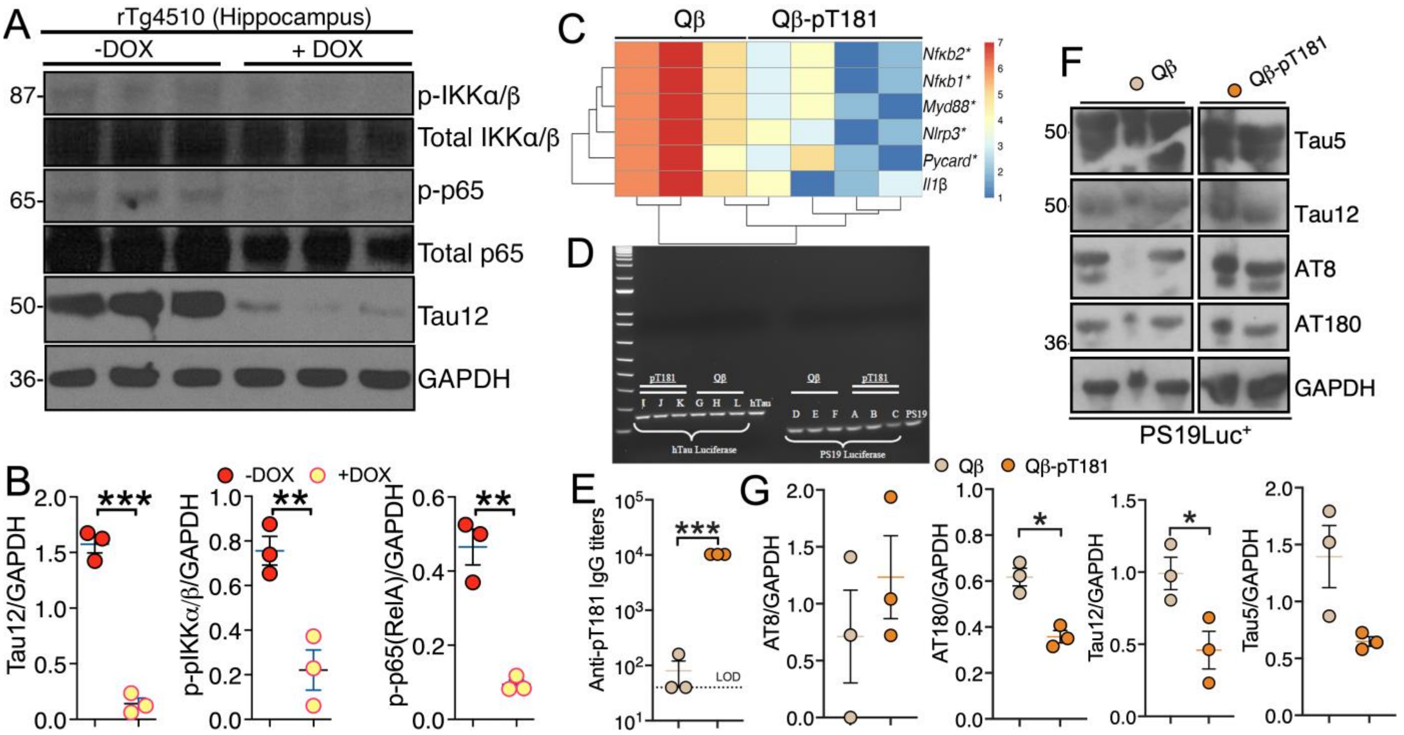

**Supplemental Figure 5: Reducing pathological tau levels chemically (via Doxycycline) or vaccines (pT181-Qβ) reduces NF-κB activation. (A, B)** rTg4510 pregnant mice were maintained in a doxycycline diet (or control diet) until the pups were born. Then the pups were also maintained on a doxycycline diet after weaning for up to 2 months. Western blot from hippocampal lysates from these up to 2-months of age doxycycline treated mice shows significantly reduced human tau (Tau12) expression, coupled with significant reduction in both total (not quantified) and phosphorylated IKKα/β and p-P65 S536 levels. **(C)** Three-month-old PS19 mice vaccinated with Qβ or pT181-Qβ, two doses and three weeks apart, show a significant reduction in the mRNA levels for the *Nfkb1*, *Nfkb2*, *Myd88*, *Nlrp3* and *Pycard* (*ASC*) and *Il1b*. **(E)** Anti-pT181 antibody titers in the sera of two-month-old albino PS19Luc+ mice vaccinated with Qb-pT181 compared to Qβ control. **(F, G)** Significant reduction in AT180/GAPDH and Tau12/GAPDH ratio in Qβ-pT181 vaccinated albino PS19Luc$^{+}$ mice compared to Qβ vaccinated control albino PS19Luc$^{+}$ mice. Data displayed as mean + SEM, unpaired t-test, *p<0.05, with n=3.

**Supplemental Table S1**

| REAGENT NAME | SOURCE | IDENTIFIER |
|---|---|---|
| **Antibodies** | | |
| Rabbit polyclonal anti-ASC | AdipoGen Life Sciences | Ca t# AG-25B-0006-C100; RRID: AB_2885200 |
| Mouse monoclonal phospho-T231 tau (AT180) | ThermoFisher | Cat # MN1040; RRID: AB_223649 |
| Mouse monoclonal phospho-S202 tau (AT8) | ThermoFisher | Cat # MN1020; RRID: AB_223647 |
| Rabbit polyclonal anti-Caspase-1 | Santa Cruz Biotech | Cat # sc-514; RRID: AB_2068895 |
| Mouse monoclonal anti-GAPDH | Millipore | Cat # CB1001-500UG; RRID: AB_2107426 |
| Mouse monoclonal anti-GAPDH | Proteintech | Cat # 6004-1-Ig |
| Rabbit monoclonal anti-β-Actin | Cell Signaling | Cat # 4970s |
| Rabbit monoclonal anti- β-Actin | ABclonal | Cat # AC026 |
| Rabbit monoclonal anti-Iba1 | Wako | Cat# 019-19741; RRID: AB_839504 |
| Goat polyclonal anti-IL-1β | R&D System | Cat # AF-401-NA; RRID: AB_416684 |
| Mouse monoclonal anti-phospho-S396/S404 tau (PHF1) | Gift from Dr. Peter Davies | [1] |
| Rabbit monoclonal anti-phospho-NF-κB p65 S536 | Cell Signaling | Cat # 3033s |
| Mouse monoclonal anti-Tau5 | ThermoFisher | Cat # AHB0042; RRID: AB_1502093 |
| Mouse monoclonal anti-Tau12 | Millipore | Cat # MAB2241; RRID: AB_1977340 |
| Mouse monoclonal anti-IκBα | Cell Signaling | Cat # 4814 |
| Mouse monoclonal anti-phospho-IκBα S32/36 | Cell Signaling | Cat # 9246 |
| Mouse monoclonal anti-NF-κB p65 | Cell Signaling | Cat # 6956 |

| | | |
|---|---|---|
| Rabbit monoclonal anti-phospho-NF-κB p65 S536 | Cell Signaling | Cat # 3033s |
| Rabbit polyclonal anti-IKKα | Cell Signaling | Cat # 2682 |
| Rabbit monoclonal anti-IKKβ | Cell Signaling | Cat # 2370 |
| Rabbit monoclonal anti-phospho-IKKα/β (Ser176/177) | Cell Signaling | Cat # 2078 |
| Goat anti-rabbit IgG - HRP | Jackson Immuno Research | Cat # 111-035-144; RRID: AB_2307391 |
| Donkey anti-goat IgG-HRP | Santa Cruz Biotech | Cat # sc-2020; RRID: AB_631728 |
| **Biological Samples** | | |
| Human Brain Tissues | NWNADC | N/A |
| **Chemicals, Peptides, and Recombinant Proteins** | | |
| TRIzol™ Reagent | ThermoFisher | Cat # 15596026 |
| LPS | Sigma-Aldrich | Cat# L2880-25MG |
| Cycloheximide (CHX) | Cayman | Cat # 26924 |
| Lactacystin | Cayman | Cat # 70980 |
| MG132 | Cayman | Cat # 13697 |
| Bafilomycin A1 (Baf A1) | Cayman | Cat # 11038 |
| PD 150606 | Cayman | Cat # 13859 |
| N-Laurylsarcosine sodium salt | Sigma-Aldrich | Cat # 61745-250G |
| 2-mercaptoethanol | Sigma-Aldrich | |
| 3-[(3-cholamidopropyl) dimethylammonio]-1-propanesulfonate (CHAPS) | ThermoFisher | Cat # 28300 |
| **Critical Commercial Assays** | | |
| High-Capacity cDNA Reverse Transcription Kit | ThermoFisher | Cat # 4368813 |
| TaqMan™ Gene Expression Assay (FAM) | ThermoFisher Scientific | Cat # 4352339 |
| Amaxa® Cell Line Nucleofector® Kit V | Lonza Bioscience | Cat # VCA-1003 |
| RNeasy Mini Kit | Qiagen | Cat # 74104 |
| **Experimental Models: Cell Lines** | | |
| C20 cell line | Gift from Dr. David Alvarez-Carbonell | Garcia-Mesa et al, 2017 |
| BV2 cell line | Gift from Dr. Gary Landreth | RRID: CVCL_0182 |
| Mouse bone marrow macrophages (BMMs) | N/A | Prepared in house |
| **Experimental Models: Organisms/Strains** | | |

| | | |
|---|---|---|
| C57BL/6J mice | Jackson Laboratory | Cat # 000664; RRID: IMSR_JAX:000664 |
| PS19 mice | Jackson Laboratory | Yoshiyama et al., 2007 |
| hTau mice | Generated previously | [2-4] |
| NFκB-GFP-Luciferase (NGL or Luc$^+$) mice | In this paper | Everhart et al, 2006 |
| hTauLuc$^+$ mice | In this paper | N/A |
| PS19Luc$^+$ mice | In this paper | N/A |
| *Mapt*$^{-/-}$ mice | Jackson Laboratory | Dawson et al., 2001; Maphis et al., 2017 |
| **Software and Algorithms** | | |
| ImageJ | [5] | https://imagej.net/; RRID:SCR_003070 |
| AlphaEaseFC™ | AlphaInnotech | http://genetictechnologiesinc.com/alpha/alpha_ease_fc.htm |
| Living Image | IVIS Lumina K series | https://resources.perkinelmer.com/lab-solutions/resources/docs/bro_010789c_01%20prd_ivis_lu minak.pdf |
| Adobe Photoshop CC | Adobe | https://www.adobe.com/products/photoshop. html; RRID: SCR_014199 |
| Prism | GraphPad | https://www.graphpad.com/scientific- software/prism/; RRID: SCR_002798 |
| **Other** | | |
| VECTASHIELD Mounting Medium with DAPI | Vector Labs | Cat # H1200 |
| Protease inhibitor cocktail | Sigma-Aldrich | Cat # P8340 |
| Phosphatase inhibitor cocktail | Sigma-Aldrich | Cat # P5726 |
| PhosSTOP® phosphatase inhibitor cocktail Tablet | Roche | Cat # 4906845001 |
| PMSF | Sigma-Aldrich | Cat # 93482-50ML-F |
| RIPA buffer | ThermoFisher | Cat # 89901 |
| Tissue protein extraction reagent | ThermoFisher | Cat # 78510 |

| | | |
|---|---|---|
| Lithium dodecyl sulfate (LDS) | ThermoFisher | Cat # B0007 |
| Reducing Agent | ThermoFisher | Cat # NP0009 |
| NuPAGE™ 4-12% Bis-Tris Protein Gels | | Cat # NP0335BOX |
| PVDF Transfer Membranes, 0.2µm | ThermoFisher | Cat # 88520 |
| ECL substrate | ThermoFisher | Cat # 34577 |
| SuperSignal™ West Pico PLUS ECL substrate | ThermoFisher | Cat # 34577 |
| Prestained protein ladder | ThermoFisher | Cat # 26616 |
| Bovine serum albumin | Sigma | Cat # 9647 |
| Blotting grade blocker nonfat dry milk | Bio-Rad | Cat # 1706404XTU |
| Percoll | GE Life Sciences | Cat # 17089102 |
| Qβ virus-like particle | Previously described | (Maphis et al., 2019) |
| Qβ-pT181 virus-like particle | Previously described | (Maphis et al., 2019) |
| **PCR primer pairs** | | |
| P301S F: 5’- CTA GAC CAC GAG AAT GCG AAG-3’ | | |
| P301S R: 5’- CTT TTG TCA CTC GGC TTT GG-3’ | | |
| hTau F: 5’- TCG TGA CCA CCC TGA CCT AC-3’ | | |
| hTau R: 5’- CGT TGT GGC TGT AGT AGT TG-3’ | | |
| mTau F: 5’- CTG CTC CAA GAC CAA GAA GGA-3’ | | |
| mTau R: 5’- TGT GTA TGT CCA CCC CAC TGA-3’ | | |
| Luc F: 5’- CCG CTT AAC AGC GTC AAC A-3’ | | |
| Luc R: 5’- GAA CTT CAG GGT CAG CTT GC-3’ | | |
| *Mapt*$^{-/-}$: F: 5’-GCC AGA GGC CAC TTG TGT AG-3’ | | |
| *Mapt*$^{-/-}$: R: 5’-ATT CAA CCC CCT CGA ATT TT-3’ | | |
